# Three-dimensional genome architecture connects chromatin structure and function in a major wheat pathogen

**DOI:** 10.1101/2025.05.13.653796

**Authors:** Ivona Glavincheska, Cecile Lorrain

## Abstract

**Background:** Genome spatial organization plays a fundamental role in biological function across all domains of life. While the principles of nuclear architecture have been well-characterized in animals and plants, their functional relevance in filamentous fungi remains largely uncharacterized. The wheat pathogen *Zymoseptoria tritici* presents a unique model for genome evolution, with a compartmentalized genome comprising conserved core and highly variable accessory chromosomes linked to genome plasticity. Here, we present the first 3D genome analysis of a eukaryotic organism with an extensive set of accessory chromosomes, revealing a hierarchical genome architecture integrating core and accessory regions.

**Results:** At the nuclear level, centromere clustering defines the global genome conformation. Accessory chromosomes are spatially segregated from core arms but maintain focal contacts with pericentromeric regions of core chromosomes, contributing to mitotic stability. At finer resolution, we identify homotypic interactions among heterochromatin-rich compartments and self-interacting domains demarcated by specific histone marks, gene expression profiles, and insulator-like sequence motifs. Notably, a subset of highly insulated, transposon-rich heterochromatic domains forms strong inter-domain interactions. Additionally, domains defined under axenic conditions with coordinated transcriptional activation during wheat infection suggest a link between 3D architecture and dynamic gene regulation.

**Conclusion:** Our study uncovers the multi-scale principles of nuclear organization in a major fungal plant pathogen and reveals how hierarchical nuclear architecture contributes to gene expression coordination and genome stability. These findings establish a conceptual framework for investigating 3D genome function and chromatin-mediated regulation in filamentous fungi and other eukaryotic microbes.

## BACKGROUND

The spatial organization of eukaryotic genomes is fundamental to cellular function, underpinning chromosomal dynamics, and contributes to genome integrity and regulation [1–3]. The physical arrangement of the chromatin in the nuclear space reflects a highly ordered and dynamic hierarchical architecture. Advances in chromosome conformation capture integrated with high-throughput sequencing (Hi-C) have significantly transformed our understanding of genome organization. In addition, Hi-C combined with polymer physics models, can be used to differentiate between chromatin states and describe chromosomal conformation [4]. The comparison of 3D genome organization across the tree of life highlights both shared features and distinct aspects of genome organization between organisms, raising the question of whether common principles govern genome architecture [5].

At the largest scale, chromosomes are organized into distinct conformations within the nucleus, generally categorized into two main types: chromosome territory-like and Rabl-like organizations [2]. Chromosome territory-like organization, primarily observed in mammals, is characterized by minimal inter-chromosomal interactions, with each chromosome occupying a discrete nuclear region during interphase [2]. In contrast, Rabl-like configurations are characterized by one or more of the following features: inter-centromere clustering, inter-telomere clustering, or parallel chromosomal alignment with both centromere and telomere clustering [2]. These features are reminiscent of the classic Rabl configuration first described in mitotic cells, where centromeres and chromosome arms are arranged in parallel [6]. The distinction between territory-based and Rabl-like chromosome conformations is thought to be driven by the presence or absence of the condensin II complex [2]. Beyond these large-scale arrangements, chromatin is further organized into epigenetically distinct domains known as A and B compartments, which preferentially interact with regions of similar chromatin state [3,5,7]. In metazoan genomes, A compartments typically correspond to gene-rich, euchromatic regions associated with active transcription, while B compartments are enriched in heterochromatic marks, depleted in genes, and generally transcriptionally repressed. This bipartite organization reflects a fundamental level of genome partitioning into functionally distinct chromatin states. In Hi-C data, these compartments are typically inferred from principal component analysis (PCA) of the contact matrix, where the first eigenvector separates regions with preferential self-interaction into A- or B-type compartments based on their chromatin and transcriptional features [8,9]. At a finer resolution, chromatin is organized into topologically associating domains (TADs), self-interacting regions within which DNA sequences interact more frequently than with sequences outside the domain [8]. The function of TADs remains debated, but they have been described as playing an important role in regulating gene expression, partly by modulating the interactions between regulatory elements and their target genes [11–13]. TAD boundaries act as insulators that constrain or mediate chromatin loop extrusion to ensure proper genomic interactions. Boundaries are enriched for architectural proteins, such as the cohesin complex and the DNA-binding protein CCCTC-binding factor (CTCF), and frequently coincide with regions containing highly transcribed housekeeping genes [11]. The disruption of genome folding can disrupt interactions between genes and regulatory elements, as observed in the enhancer hijacking of proto-oncogenes [12,13]. In contrast to metazoan systems, other organisms exhibit 3D genome organization patterns that, although reminiscent of features such as loops and compartments observed in mammals, arise through distinct mechanisms. In plants, chromatin is arranged into self-interacting domains associated with different transcriptional and epigenetic states, such as gene-active, Polycomb-repressed, and intermediate regions [8,17]. Although Hi-C maps of plant genomes reveal features resembling metazoan TADs, with boundaries often enriched in highly transcribed or epigenetically marked loci, plants lack orthologues of canonical architectural proteins like CTCF [8,14–16]. Furthermore, in large plant genomes, genes separated by heterochromatic regions frequently engage in long-range looping, suggesting a genome organization driven more by structural constraints than by enhancer-promoter interactions [14,17]. Beyond eukaryotes, Hi-C experiments in prokaryotes have revealed that bacterial and archaeal genomes are also spatially structured. In bacteria, chromosomal interaction domains (CIDs) represent nested, self-associating units that are typically demarcated by highly expressed genes [18–20]. CIDs reflect an organization principle where operon-level co-regulation may be shaped by spatial proximity. In archaea, more complex folding has been observed, including compartment-like patterns and transcriptionally linked CIDs, with boundary formation influenced by both SMC proteins and transcriptional activity [21,22]. These findings depict hierarchical 3D genome organization as a universal feature, but its underlying mechanisms and biological roles differ substantially across the tree of life. Understanding the establishment and regulatory impact of 3D genome organization is therefore essential. Yet, despite significant progress in metazoan models, these processes remain poorly understood in many other eukaryotes, particularly in fungi.

The fungal kingdom, comprising millions of species, has significant ecological and economic impacts [23]. Fungal genomes display an extraordinary range of sizes and architectures, from compact, gene-dense assemblies of under 10 Mb with very few repetitive elements, such as in *Saccharomyces cerevisiae* [24], to massive genomes exceeding 1 Gb that are dominated by transposable elements (TEs), as observed in obligate biotrophic plant pathogens where TE content can reach over 90% of the genome [25]. This variation is frequently accompanied by distinct genome compartmentalization, with some species exhibiting clear separation between gene-dense, TE-poor regions and gene-poor, TE-rich regions [26,27]. In some species, this compartmentalization extends to the chromosomal level, where dispensable accessory chromosomes are often enriched in TEs and species-specific genes [27]. Despite the remarkable genomic diversity observed across the fungal kingdom, our understanding of their 3D genome organization remains limited [28]. While certain features appear to be conserved, other aspects of nuclear architecture display substantial variability across fungal lineages, underscoring the need for broader high-resolution comparative analyses. Fungal genomes generally exhibit a Rabl-like organization characterized by centromere clustering, as demonstrated in yeast and filamentous fungi [28,29]. However, the spatial organization of accessory chromosomes remains completely unknown. Whether these chromosomes adopt similar spatial constraints as core chromosomes or are sequestered in distinct nuclear territories is currently unresolved. In addition, preferential spatial interactions among transcriptionally A and B chromatin are not uniformly observed in fungi. To date, clear evidence for A/B compartmentalization has been reported only in a limited number of fungal species, such as *Rhizophagus irregularis* [30]. Self-interacting chromatin domains have been described in multiple fungal species using Hi-C approaches. While these structures resemble TADs of metazoans or CIDs of prokaryotes, their defining features vary across fungal species. In *Saccharomyces cerevisiae*, high-resolution maps revealed the presence of small self-associating domains, typically covering one to five genes [31]. These domains are delimited by nucleosome-depleted regions, often located at the promoters of highly transcribed genes, and are enriched in chromatin remodelers and transcription factors, rather than architectural proteins [45,46]. In *S. cerevisiae*, CIDs are implicated in critical genome functions such as recombination repression, genome evolution, and the temporal organization of replication origins during mitosis [31]. In the filamentous plant-pathogen *Verticillium dahliae*, larger domains identified as CIDs exhibit strong internal interactions and are flanked by regions marked with facultative heterochromatin [32]. In *Neurospora crassa*, domain-like structures consist of weakly insulated euchromatic regions interspersed with constitutive heterochromatin blocks [33,34]. These findings collectively suggest that self-interacting domains are widespread in fungi, but their size, boundary determinants, and chromatin context differ markedly between species.

The fungal wheat pathogen *Zymoseptoria tritici* emerged alongside the domestication of wheat in the Fertile Crescent [35]. The genome of *Z. tritici* is compartmentalized into thirteen core and eight accessory chromosomes [27,36]. The core chromosomes, which are gene-rich and transposable element (TE)-poor, contrast with the gene-poor and TE-rich accessory chromosomes, an ancestral feature shared across the *Zymoseptoria* genus [27]. The accessory chromosomes exhibit a high degree of presence/absence polymorphism in *Z. tritici* populations [37], and some chromosomes appear to be more prone to destabilization than others, as indicated by targeted loss experiments [38]. The genomic partitioning of *Z. tritici* is further reflected by distinct epigenetic signatures, particularly in the presence of specific histone modifications in accessory chromosomes [39]. The epigenetic landscape of *Z. tritici* exhibits the distinction of euchromatin, constitutive and facultative heterochromatin, cytosine methylation (5mC) regions, small acrocentric centromeres, and telomeric repeats, underlining the complexity of *Z. tritici*’s chromatin organization [27,40–45]. Cytological observations using GFP-tagged centromeres previously reported the presence of 4-7 centromeric foci during interphase, suggesting partial centromere clustering [39]. Pathogenicity-related genes in *Z. tritici* are enriched within TE-rich regions of the genome, which likely facilitates rapid evolution through genome rearrangements and epigenetic regulation [45,46]. These genes are predominantly repressed in the absence of the host and maintained in a transcriptionally silent state by heterochromatin marks [46]. Upon host colonization, chromatin remodeling events activate the expression of pathogenicity-related genes [46]. Despite these detailed insights into the genomic and epigenomic features of *Z. tritici*, the three-dimensional organization of its genome remains uncharacterized. In particular, how chromatin domains are spatially arranged, whether accessory chromosomes occupy distinct nuclear territories, and the extent to which point centromeres cluster within the nucleus are open questions in the field. Furthermore, the presence and determinants of self-interacting chromatin domains in *Z. tritici* have yet to be investigated. This study addresses these gaps by integrating high-resolution Hi-C data with genomic, epigenomic, and transcriptomic information to comprehensively characterize the 3D genome architecture of *Z. tritici*. To our knowledge, this represents the first in-depth analysis of spatial genome organization in a eukaryotic microbe with a compartmentalized genome that includes eight accessory chromosomes. By mapping the folding principles of core versus accessory chromosomes, assessing centromere organization, and identifying chromatin domains, our work provides new insights into the interplay between genome structure, epigenetic regulation, and adaptive potential in fungal pathogens.

## RESULTS

### The 3D genome of *Z. tritici* reveals distinct conformation patterns between core and accessory chromosomes

To investigate the three-dimensional (3D) genome organization of *Z. tritici* IPO323, we generated high-resolution Hi-C sequencing revealing a Rabl-like architecture and previously undescribed spatial segregation between core and accessory chromosomes. We performed Hi-C on two independent replicates which showed high reproducibility (Pearson correlation = 0.87; 77–81% valid read pairs; Supplementary Fig. S1). After merging replicates, we obtained ∼19.2 million deduplicated valid read pairs (Supplementary Table S1), enabling chromatin interaction detection at a minimal bin size of 3 kb. At this resolution, 80% of bins on core chromosomes were supported by >1,000 contacts, with nearly complete coverage at 10 kb on all chromosomes except chromosome 21 (Fig. 1A; Supplementary Table S2). As in other fungi, local intrachromosomal contacts dominated the Hi-C map, yielding a strong diagonal (Fig. 1A). The strongest interchromosomal contacts were enriched at centromeres and pericentromeric regions, consistent with a Rabl-like conformation (observed-to-expected contact > 3.5; Fig. 1A, 1B; Supplementary Fig. S2-S3). These centromeric regions were previously mapped in IPO323 using CENH3 (CENP-A homolog) ChIP-seq [39] and their enrichment in inter-chromosomal contacts independently confirms their spatial colocalization in the nucleus, as expected under a Rabl-like organization. Outside of centromere-centromere interactions, strong interchromosomal contacts were also enriched between regions marked by constitutive heterochromatin (H3K9me3; Supplementary Fig. S3; Supplementary Table S3). Notably, 32.6% of the strong interchromosomal interactions (observed-to-expected contact > 3.5) occurred between two H3K9me3-enriched bins, revealing a heterochromatin network spanning multiple chromosomes (Supplementary Fig. S3C-D).

**Fig. 1:**
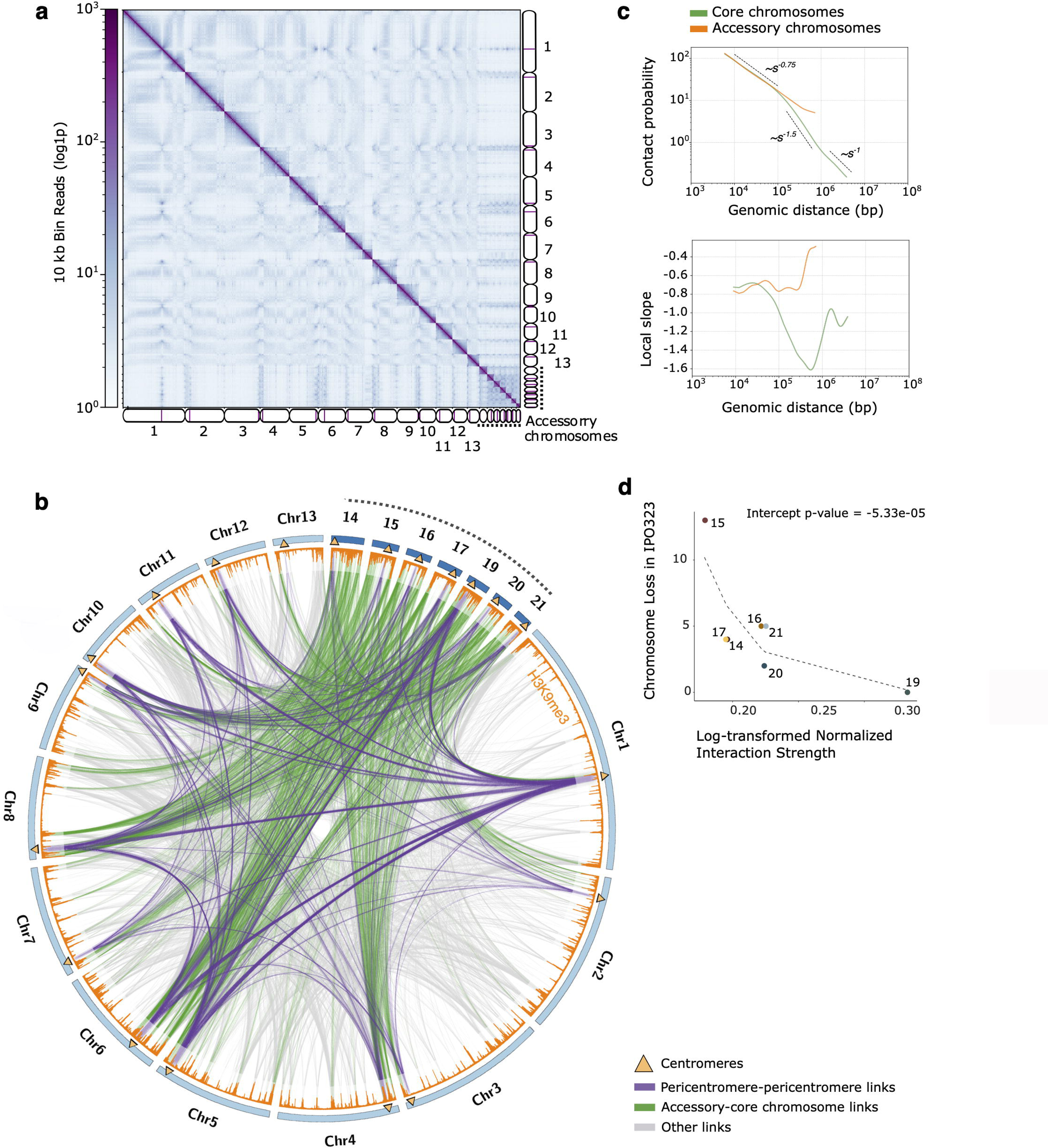
The Rabl-like three-dimensional structure of *Zymoseptoria tritici* is compartmentalized. a) Heatmap of genome-wide normalized Hi-C contact map at 10kb resolution. Contact numbers per 10kb bins are represented in a log scale. Chromosomes are represented by ideograms with purple marking the centromeres [39]. b) Strongest inter-chromosomal interactions with an observed-to-expected value >3.5 at 10 kb bin resolution. Core chromosomes are represented in light blue. Accessory chromosomes are represented in dark blue. Centromeres are marked by triangles. H3K9me3 enrichment is shown in the orange histogram. Links represent inter-chromosomal pericentromere-pericentromere interactions (purple), accessory-core chromosome (green), or any other interactions (grey). c) Contact probability (top) and contact frequency decay slope (bottom) against genomic distance for core (green) and accessory (orange) chromosomes. d) Relationship between chromosomal interaction strength and chromosome loss in *Zymoseptoria tritici* strain IPO323, after carbendazim treatment [38]. Scatterplot showing the association between log-transformed normalized Hi-C interaction strength (x-axis) and the frequency of chromosome loss (y-axis) across individual chromosomes. A dashed grey line indicates the predicted linear relationship. The model indicates a significant inverse relationship between interaction strength and chromosome loss (Poisson GLM, p-value < 0.05).

Accessory chromosomes occupy a defined spatial niche within the nucleus, exhibiting preferential associations with the pericentromeric regions of core chromosomes. Specifically, 78.6% of core-accessory interactions involved contacts spanning the full length of accessory chromosomes but were restricted to the pericentromeric regions of core chromosomes (observed-to-expected contact > 3.5; Fig. 1B; Supplementary Fig. S3C). This spatial organization suggests that accessory chromosomes are tethered to core chromosomes pericentromeric regions, while mostly remaining separated from core chromosome long arms (Fig. 1B). These findings provide the first spatial insight into accessory chromosome positioning in a fungal genome and support a model in which centromere-proximal anchoring contributes to higher-order genome compartmentalization. Given the known instability of accessory chromosomes [38], we tested whether their nuclear interactions influence their maintenance. Using log-transformed Poisson regression, we found a significant negative association between interaction strength (normalized for accessory chromosome size) and chromosome loss frequency following β-tubulin inhibition in IPO323 [38] (p-value = 5.33e-05 for the intercept; p-value = 0.0025 for normalized interaction strength; Fig. 1D). This result supports a model in which stronger core-accessory chromosome interactions correlate with accessory chromosome stability.

To determine whether core and accessory chromosomes differ in their folding behavior, we estimated the contact frequency decay exponent (α) as a function of genomic distance. The core chromosomes exhibited an average decay exponent of α = −1.22 (standard deviation (SD) = 0.1), with three distinct scaling regimes depending on genomic separation, consistent with hierarchical chromatin organization (Fig. 1C; Supplementary Table S4). At short distances (< 70 kb), contact frequency decayed more gradually than the global average with α = −0.71 (SD = 0.01). Interactions between 70 kb and 1 Mb followed a more heterogeneous and steeper decline with α = −1.43, SD = 0.09). At distances beyond 1 Mb, contact frequency declined at a slower rate relative to intermediate distances (α = −1; SD = 0.09; Fig. 1C; Supplementary Table S4). This triphasic decay pattern reflects hierarchical chromatin folding, suggestive of domains or compartments on the core chromosomes. In contrast, the accessory chromosomes showed low difference between short- and long-distance contact frequency decay (α = −0.71 vs. α = −0.62), consistent with weak or absent domain-level organization (Fig. 1C; Supplementary Table S4). This pattern suggests that, compared with the core chromosomes, accessory chromosomes behave as unitary compartments exhibiting weak scale-dependent folding (Fig. 1C; Supplementary Table S4).

Taken together, these results reveal fundamental differences in chromatin conformation between core and accessory chromosomes in *Z. tritici*. Core chromosomes exhibit hierarchical folding, while accessory chromosomes adopt a more uniform interaction profile, indicative of a compact structure with limited internal compartmentalization. These structural differences, coupled with accessory chromosomes preferential tethering to pericentromeric regions of core chromosomes, suggests that accessory chromosomes occupy spatially constrained territories within the nucleus potentially contributing to their differential mitotic stability.

### B chromatin compartments in *Z. tritici* engage in homotypic interactions at the intrachromosomal level

To explore how 3D chromatin organization reflects functional genome partitioning in *Z. tritici*, we performed eigenvector decomposition of inter- and intrachromosomal Hi-C contact maps (Fig. 2; Supplementary Figs. S4-S5). At the interchromosomal level, the first eigenvector separated chromosome arms, with sub-telomeric regions on both arms displaying similar profiles, indicative of Rabl-like interchromosomal interactions (Supplementary Figs. S4-S5). At the intrachromosomal level, eigenvector profiles captured finer-scale chromatin features and more localized and compartmentalized interaction patterns (Fig. 2; Supplementary Figs. S5). Regions with negative eigenvector values exhibited strong spatial self-association, consistent with compartmental segregation. These regions were enriched for heterochromatic histone modifications, including H3K9me2 and H3K9me3, and depleted of euchromatin-associated marks such as H3K4me2 (i.e. analogous to B compartments; Fig. 2B; Supplementary Fig. S5). In contrast, positive eigenvector regions were more often associated with euchromatic features (i.e. analogous to A compartments). Comparison of contact frequencies confirmed that homotypic B-B interactions were predominant, while A-A interactions were comparatively weaker (Fig. 2A). This asymmetry indicates that compartmentalization in *Z. tritici* is primarily driven by interactions among transcriptionally silent, heterochromatic domains, rather than by the formation of discrete euchromatic A compartments.

**Fig. 2:**
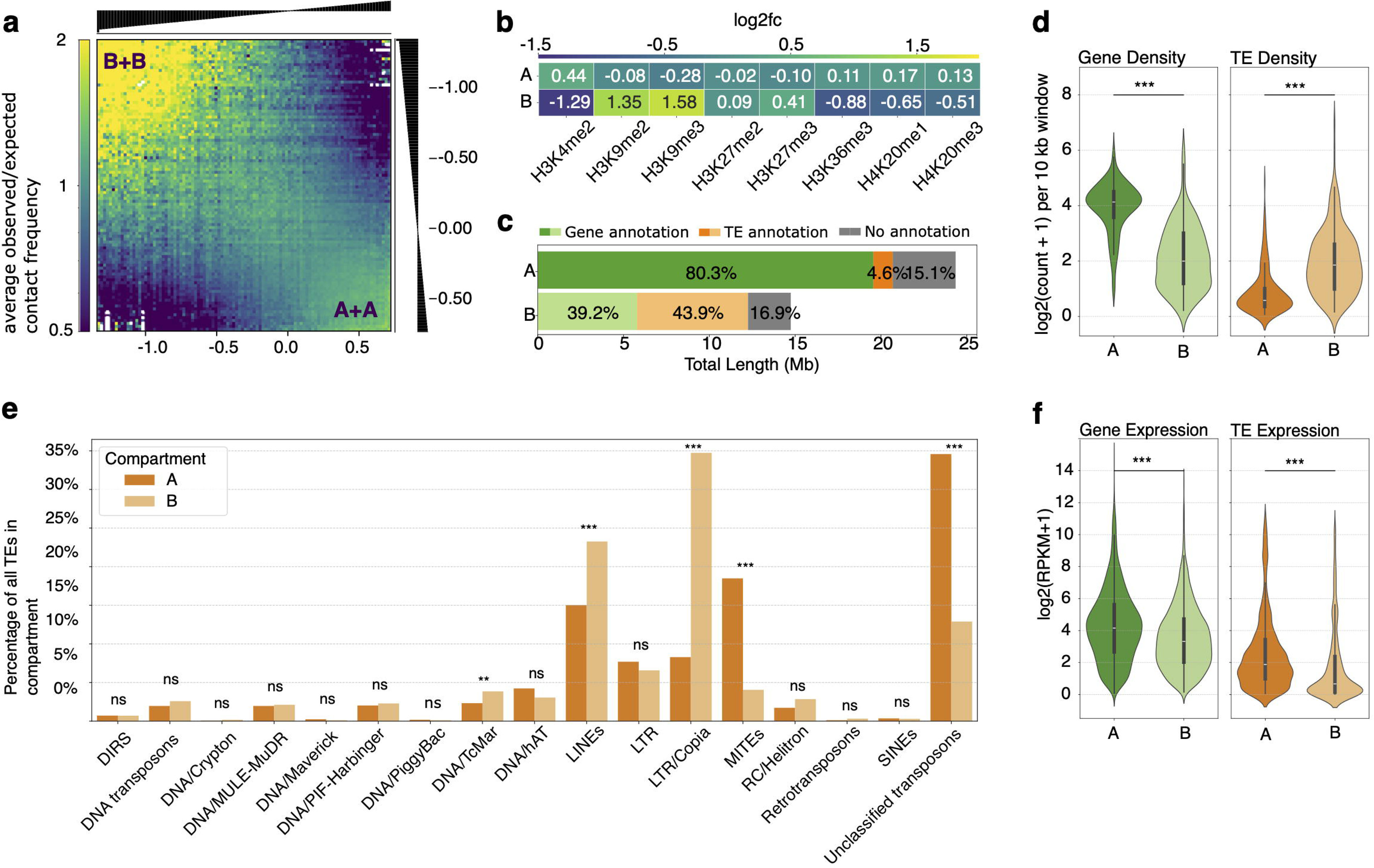
Homotypic interactions and epigenomic features of A/B compartments in the *Zymoseptoria tritici* genome. a) Saddle plot representing genome-wide intrachromosomal Hi-C interactions at 3 kb resolution. Homotypic B-B interactions are represented in the top-left, while A-A interactions are observed in the bottom-right. b) Enrichment of histone modifications across A and B compartments, highlighting distinct chromatin states. c) Genomic composition of A and B compartments, showing proportions of genes (green), transposable elements (orange), and intergenic regions (grey). d) Density of genes (green) and transposable elements (orange) across A and B compartments. e) Distribution of transposable element families within A and B compartments. Asterisks indicate statistically significant differences in TE content based on pairwise Fisher exact tests with FDR correction using the Benjamin-Hochberg method (ns: non-significant; *: p-value < 0.05; **: p-value < 0.01; ***: p-value < 0.001). f) *In vitro* Expression levels of genes (green) and transposable elements (orange) in A and B compartments.

The B-compartment blocks had an average size of ∼60 kb, accounting for 14.74 Mb (∼36%) of the genome, and were characterized by low gene density and a high density of TEs (Fig. 2C-D, Supplementary Table S6). Notably, we observed a significant enrichment of retrotransposons, particularly LINEs and LTR elements from the Copia superfamily (Fischer exact test BH-adjusted p-value < 0.001, Fig. 2C-E, Supplementary Table S7). In contrast, the A-compartment blocks had an average length of 100 kb, spanning a total of 24.35 Mb, of which 80.3% are covered by genes (9,996 genes; Fig. 2C). The A compartments exhibited high gene density, low TE density (4.3% of TEs), and enrichment in miniature inverted-repeat transposable elements (MITEs, Fischer exact test BH-adjusted p-value < 0.001; Fig. 2C-E). Gene ontology (GO) enrichment analysis in the A-compartments revealed an overrepresentation in core cellular processes including translation, ribosome function, RNA binding, and intracellular non-membrane-bounded organelles (FDR-adjusted p-value < 0.01; Supplementary Table S8-S9). Consistent with these observations, genes and TEs within A-compartment blocks exhibited significantly higher *in vitro* expression than those in B-compartment blocks (Fig. 2F), indicating that transcriptional regulation of both genes and TEs is dependent on chromatin compartmentalization.

Taken together, these findings demonstrate that the genome of *Z. tritici* is partitioned into spatially segregated chromatin compartments, with heterochromatic B compartments driving the overall compartmentalization landscape through strong intra-compartment interactions. This compartmental organization operates at the intrachromosomal level and is distinct from the global partitioning into core and accessory chromosomes, underscoring the multi-layered nature of 3D genome folding in this fungal pathogen.

### The presence of chromatin interacting domains in *Z. tritici* reveals boundaries enriched in facultative heterochromatin and specific DNA binding motifs

To determine whether the *Z. tritici* genome is organized into chromatin-interacting domains (CIDs), we calculated insulation scores from Hi-C contact maps at 3 kb resolution. We identified 513 CIDs and 533 CID boundaries in *Z. tritici* IPO323, with an average CID length of approximately 70 kb (Supplementary Table S10-12). The distribution of CIDs varied between core and accessory chromosomes, with 92% of CIDs (473 CIDs) located on core chromosomes and only 8% (40 CIDs) on accessory chromosomes (Supplementary Table S12). On average, CIDs on core chromosomes contained 25 genes and 6 TEs, whereas CIDs on accessory chromosomes contained an average of 14 genes and 17 TEs per CID (Supplementary Table S12).

Since CID boundaries act as insulators between domains, we identified them by detecting local insulation minima and further examined whether CID boundaries exhibited specific epigenomic and genomic features (Fig. 3A-B). We observed that CID boundaries were depleted of the constitutive heterochromatin histone modifications H3K9me2 and H3K9me3 but enriched for facultative heterochromatin modifications H3K36me3, H4K20me1, and H4K20me3 (Fig. 3B, Supplementary Fig. S6). In accessory chromosomes, CID boundaries were also enriched for accessory chromosome-associated modifications, H3K27me2 and H3K27me3 (Supplementary Fig. S6). In addition, CID boundaries displayed lower TE density, reduced RIP-like mutational signatures, and higher gene density compared to domain interiors. Despite lacking enrichment in the active transcription mark H3K4me2, genes and TEs at CID boundaries, exhibited comparable expression levels to those within CIDs, and reduced variability across conditions, consistent with constitutive, stable expression patterns rather than dynamic regulation (Supplementary Fig. S7). To assess whether CID boundaries in *Z. tritici* are preferentially associated with conserved or functionally specialized genes, we characterized the 613 genes overlapping (>50%) with boundary regions. Comparative analysis across 19 *Z. tritici* reference genomes [47] revealed that only 69 of these genes were conserved across all strains indicating no significant enrichment for core genes (Fisher’s test, p-value = 0.76; Supplementary Tables S13-S15). Similarly, we did not find enrichment of specific Gene Ontology (GO) terms or functional domains (Supplementary Tables S13-S15). Finally, we searched whether specific DNA binding motifs were present at CID boundaries. Motif discovery analysis uncovered 35 significantly enriched known and *de novo* DNA binding motifs, which clustered into nine motif groups (Supplementary Table S16). Among these, one of the strongest matches showed similarity to the consensus binding motif of the mammalian KRAB zinc finger protein ZNF263, a factor recently implicated in establishing insulator activity at chromatin domain boundaries [48]. IPO323 has a single protein annotated with a KRAB-like zinc finger domain (ZtIPO323_082310), but this protein exhibits low sequence similarity in its zinc finger domains with the human ZNF263 protein, suggesting potential convergent motif recognition or functional analogs with distinct primary sequences. (Supplementary Table S17).

**Fig. 3:**
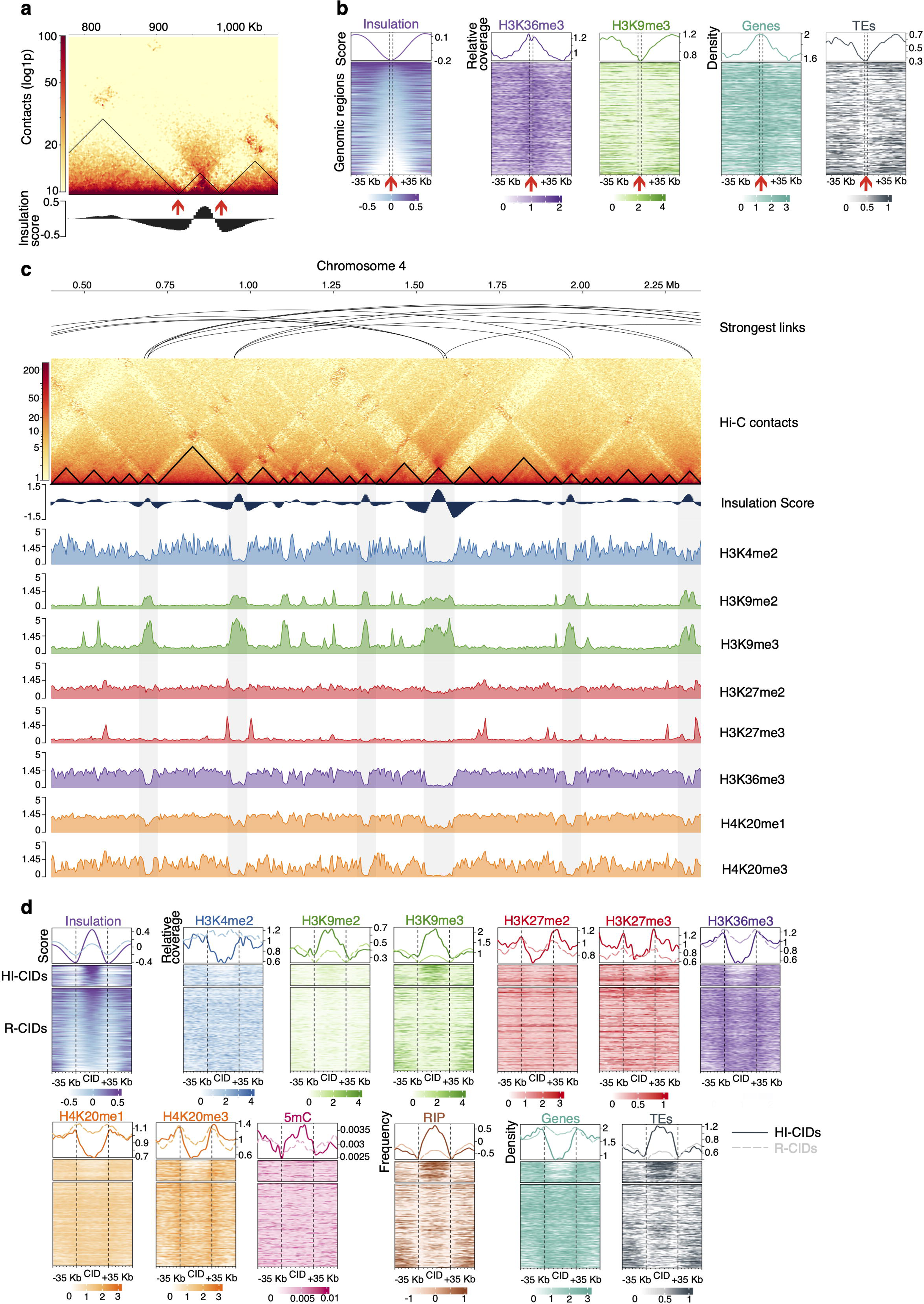
The genome of *Zymoseptoria tritici* forms chromatin interacting domains (CIDs) with specific genomic and epigenomic signatures. a) Close-up of Hi-C contact map showing local interaction frequencies at 3 kb bins resolution and detection of self-interacting CID structures. Insulation score was calculated with a genome-wide sliding window approach. Predicted CIDs are represented by black triangles, and boundaries are indicated by red arrows. b) Heatmap of chromatin insulation scores, histone modifications (H3K36me3 and H3K9me3), gene, and TE density enrichment centered on domain boundaries, with 35 kb upstream and downstream regions shown per row. The top panel shows the average profile across all boundaries depicted in the heatmap. c) Close-up example of CID annotations on chromosome 4 of *Z. tritici* IPO323. From top to bottom: strongest intra-chromosomal links (observed/expected > 1.75) across chromosome 4; chromosome 4 Hi-C contact map with CID annotated by triangles; insulation scores within chromosome 4; histone modifications for euchromatin H3K4me2, heterochromatin H3K9me2, H3K9me3, and facultative heterochromatin H3K27me2, H3K27me3, H3K36me3, H3K20me1, H4K20me3. d) Heatmap of chromatin insulation scores, histone modifications, 5-methylcytosine DNA methylation (5mC), repeat-induced point mutation (RIP) composite index frequency, genes, and TEs density centered on CIDs, with 35 kb upstream and downstream regions shown per row. The top panel shows the average profile across all CIDs depicted in the heatmap. Highly-insulated CIDs (HI-CIDs) are depicted by solid lines, and “regular-CIDs” (R-CIDs) are depicted by dashed lines.

Together, these results suggest that while CID boundaries in *Z. tritici* are not enriched in conserved or functionally specialized genes, they exhibit distinct epigenetic and sequence signatures consistent with a potential architectural role in genome organization. The enrichment of ZNF263-like motifs points to a potential mechanism of boundary insulation mediated by sequence-specific binding, possibly involving fungal architectural proteins.

### Highly insulated CIDs define chromosomal conformation, while local CID organization regulates transcriptional dynamics during plant infection

We next investigated the internal organization of CIDs to evaluate their contribution to higher-order chromatin conformation and transcriptional regulation. We observed substantial heterogeneity in intrachromosomal contact patterns across CIDs, with a subset displaying pronounced insulation from flanking regions, hallmarks of locally compartmentalized chromatin architecture (Fig. 3C-D). To systematically identify these highly insulated domains, we compared maximum insulation scores within each CID to those of their flanking boundaries. CIDs with insulation strength difference exceeding the genome-wide median were classified as highly insulated CIDs (HI-CIDs), while the remainder were designated as regular CIDs (R-CIDs). This analysis yielded 105 HI-CIDs, representing 20% of all CIDs genome-wide (Supplementary Table S18). Analysis of strongest interactions (observed-to-expected intrachromosomal contacts > 1.75) revealed significantly higher interaction frequencies between HI-CIDs compared to R-CID/R-CID and HI-CID/R-CID combinations (Mann-Whitney p-value < 0.05; Supplementary Fig. S8), indicating that HI-CIDs form a distinct interaction network within the genome. In addition, HI-CIDs were characterized by enrichment in heterochromatin-associated histone marks (notably H3K9me2 and H3K9me3), and a relative depletion of active and facultative chromatin modifications (Fig. 3C-D). Consistent with their chromatin profile, HI-CIDs exhibited elevated TE density, reduced gene content, and low transcriptional activity under *in vitro* conditions (Supplementary Fig. S9). Notably, HI-CIDs were also enriched in predicted effector genes, implying that heterochromatin dynamics within these regions may be restructured to enable effector gene activation during host infection (Fisher’s exact p-value < 0.01; odds ratio = 1.54; Supplementary Tables S19).

To investigate whether CID organization could influence co-expression patterns in *Z. tritici*, we analyzed transcriptomic data obtained from both axenic culture and four stages of wheat infection. We specifically asked whether genes and TEs located within the same CID (i.e. defined based on Hi-C data from axenic culture) exhibit coordinated expression changes across these distinct conditions. Using a linear model, we tested whether CID membership could explain patterns of differential expression between *in vitro* and *in planta* conditions (Supplementary Tables S20–S23). The model significantly outperformed a null expectation (likelihood ratio test p-value < 0.01) and explained between 5% and 9% of the adjusted variance in gene and TE expression (Supplementary Table S24), indicating that CID organization may contribute to transcriptional dynamics during infection. To further assess the relative contribution of CID organization to transcriptional variation, we extended our linear regression model to include gene GC content and distance to the nearest TE as additional predictors. Including these variables alongside CID membership modestly improved the model’s explanatory power, increasing the proportion of variance explained by an additional 3% to 6% (Supplementary Table S24). However, in the absence of CID membership as a predictor, gene GC content and TE proximity alone accounted for less than 0.1% of the observed transcriptional variance, reinforcing the importance of domain-level chromatin organization in regulating gene expression.

To identify specific CIDs exhibiting coordinated transcriptional behavior, significant CIDs from the linear regression were classified as co-regulated for CIDs containing at least 10 genes and TEs. This analysis revealed 46 co-regulated CIDs (8.9% of all CIDs) that exhibited significant coordination during at least one stage of infection (Supplementary Fig. S10; Supplementary Table S25). Notably, 23 of these CIDs had coordinated expression observed only during a single infection phase, suggesting that dynamic chromatin restructuring may contribute to phase-specific transcriptional responses. A substantial fraction of co-regulated domains localized to regions with accessory-like features, including 19 on accessory chromosomes and six on the long arm of chromosome 7, which is enriched in H3K27me3 and exhibits low gene density (Supplementary Fig. S11). Additionally, 16 of the 46 co-regulated CIDs were classified as highly insulated (HI-CIDs), further linking chromatin insulation and domain architecture to coordinated regulation. Collectively, these results show that a subset of CIDs functions as co-regulatory units in *Z. tritici*, and that heterochromatic features, insulation strength, and chromosomal context together shape transcriptional coordination during host infection.

## DISCUSSION

Our study provides the first comprehensive characterization of the hierarchical 3D chromatin organization in the major fungal wheat pathogen *Z. tritici*. The genome of *Z. tritici* adopts a Rabl-like nuclear conformation, consistent with other fungal genomes [30,32,34,49,50] but with distinct folding patterns between core and accessory chromosomes. Accessory chromosomes are anchored to the pericentromeric regions of core chromosomes, with interaction strength correlating with stability, indicating that pericentromeric association may reinforce their nuclear positioning and stability. We detected hierarchical folding within core chromosomes, while accessory chromosomes displayed a more uniform organization. Beyond this global architecture, we identified intrachromosomal compartments, with B compartments engaging in homotypic interactions and enriched for retrotransposons. Furthermore, we detected self-interacting CID structures with boundaries enriched in facultative heterochromatin marks, stable gene expression and putative insulator motifs that resemble the binding sites of a mammalian architectural protein. Additionally, we defined highly insulated CIDs that form an interaction network of heterochromatic islands and harbor genes related to pathogenicity. A subset of CIDs showed transcriptional co-regulation patterns that varied across infection stages, indicating a regulatory role for CID organization in infection stage-specific gene expression. Our results establish a multilayered view of 3D genome organization in *Z. tritici*, revealing chromatin conformation as a key component of genome function.

The core-accessory genome compartmentalization of *Z. tritici* extends beyond linear genome characteristics and is mirrored in the three-dimensional organization of the nucleus. Within this nuclear context, accessory chromosomes are preferentially associated with the pericentromeric regions of core chromosomes while remaining excluded from interactions with core chromosome arms. Interestingly, the accessory chromosomes harboring the weakest interactions with core chromosomes, specifically their pericentromeric regions, were also the ones most frequently lost following treatment with carbendazim, a fungicide that disrupts β-tubulin polymerization and impairs mitotic spindle assembly [38,51]. This observation suggests that robust pericentromeric tethering may play a stabilizing role for accessory chromosomes during mitosis. Similar phenomena have been observed in *S. cerevisiae*, where microtubule-disrupting agents reduce centromere clustering, implying that centromeres are bundled at the spindle pole body during interphase [52,53]. These parallels suggest that centromeric architecture in *Z. tritici* may serve a comparable stabilizing function. Further supporting this hypothesis, accessory chromosomes with stronger pericentromeric associations, such as chromosome 19, exhibited greater stability under carbendazim treatment [38]. This chromosome also shows high prevalence in natural populations of *Z. tritici* [37], aligning spatial stability with evolutionary persistence. In contrast, despite the general presence/absence polymorphism trend among accessory chromosomes, those most frequently lost under experimental conditions were not necessarily the least prevalent in global populations [37,44]. One possible explanation for this discrepancy is that the strength of core-accessory chromosome interactions may be strain-specific. It is likely that in different natural strains of *Z. tritici*, the core-accessory interactions may vary, affecting the stability and presence of accessory chromosomes in a context-dependent manner. Additionally, evolutionary pressures and local adaptation may influence the retention or loss of these chromosomes, contributing to the observed diversity in the presence of accessory chromosomes across strains [38]. Further research, including multiple-strain Hi-C analyses, could provide insights into how core-accessory chromosome interactions have shaped the genomic architecture of *Z. tritici*.

*Z. tritici* genome displays a global Rabl-like organization. Cytological observations using GFP-tagged CENH3 further support the Rabl-like nuclear organization, revealing that *Z. tritici* centromeres cluster into 4 to 7 distinct nuclear foci per nucleus during interphase [39]. These chromocenters were observed in individual spores, indicating variability in centromere clustering from cell to cell. In contrast, our Hi-C data reflect an averaged pattern of centromere-centromere interactions across a spore population, revealing a consistent signal of clustering. The convergence of these single-cell and population-level datasets suggests that while the number and position of chromocenters may vary between nuclei, centromeres consistently associate into a limited number of foci, collectively shaping a Rabl-like nuclear architecture. Beyond this centromere-driven global organization, the Hi-C data also reveal fine-scale compartmentalization within chromosomes. Notably, the B compartments exhibit strong homotypic interactions, forming a spatially distinct subnetwork largely insulated from the euchromatic A compartments, showing minimal inter-compartment connectivity. Unlike the well-defined A/B compartmentalization observed in the human genome [3] or the checkerboard interaction patterns reported in *R. irregularis* [30], *Z. tritici* displays a genome topology that is dominated by internal B-B interactions. Compartmentalization inferred through principal component analysis of Hi-C contact matrices revealed that B compartments are small (∼60 kb), with a similar size and positioning of CIDs identified in our study. Specifically, these B compartments co-localize with highly insulated CIDs that form an interlinked network of heterochromatic regions. This suggests that *Z. tritici* chromatin exhibit a heterochromatin-driven architecture, where B compartments and highly insulated CIDs act as spatial reservoirs for transcriptionally repressed, TE-rich genomic regions. This compartmentalization likely reinforces silencing by sequestering these regions away from the transcriptionally active A compartments. Such an arrangement parallels chromatin architectures observed in other fungi [30,32,49]. For example, in *N. crassa*, heterochromatic domains marked by H3K9me3 aggregate into discrete chromatin regions that contribute to genome stability [33,54]. In *Z. tritici*, the co-occurrence of small B compartments with CID structures supports a fine-scale, heterochromatin-driven genome topology that enables both regulatory insulation and structural organization.

We identified CIDs with specific epigenetic characterization of their boundaries, supporting the presence of chromatin-interacting domains as a universal genomic feature [55]. *Z. tritici* displays CIDs of ∼70 kb in size, similar to the average CID size described in other fungi [32,49], but far smaller than metazoan TADs, which typically span over megabases [55]. The detection of CIDs in *Z. tritici* was further supported by analysis of contact probability decay slopes consistent with expectations from polymer physics modeling. These patterns, showing elevated interaction frequencies at sub-100 kb distances, are consistent with the presence of interaction domains [4]. Unlike TADs described in metazoans, we counted only a subset of co-regulated CIDs, suggesting that structural features of the genome coordinate local transcriptional regulation during host-pathogen interactions, similar to other plant-associated fungi [32,49]. In metazoans, the concept of TADs is also strongly linked to the active loop extrusion creation process, which is dependent on cohesin and the boundary-associated protein CTCF, or other insulator proteins [56]. However, self-interacting domains across the tree of life were described in the absence of insulator proteins, as well as cohesin [57,58]. For instance, CIDs in prokaryotes structure the genome at the gene operon level, with insulated boundaries associated with highly expressed genes [19,20,59]. In *Z. tritici*, CID boundaries are gene-dense, TE-poor, and enriched in facultative heterochromatin histone modifications, consistent with boundaries described in *V. dahliae* [32]. Gene expression analysis further revealed that genes at CID boundaries are not markedly different in expression level compared to those within domains, but display reduced expression variability and consistent expression across conditions. This pattern supports a model where boundary-associated genes are constitutively expressed, in line with boundary behavior in other species [32,60]. In *N. crassa*, interspersed heterochromatin blocks have been proposed to act as domain boundaries in place of insulator proteins [29,33]. However, our findings in *Z. tritici* suggest a more nuanced organization, as we observed multiple CIDs between heterochromatic blocks, with well-defined boundaries marked by distinct epigenetic profiles that are similar to those in *V. dahliae* [32]. Furthermore, we observed a significant enrichment of zinc finger transcription factor DNA-binding motifs at *Z. tritici* domain boundaries, notably those associated with the vertebrate insulator protein ZNF263 [61]. In *V. dahliae*, boundary-enriched GAAG-motifs and TATA-motifs associate with a decrease in chromatin accessibility [32]. The presence of a zinc-finger binding motifs in *Z. tritici* CID boundaries suggests that zinc-finger proteins may contribute to domain boundary formation in fungi. As a diverse and underexplored protein family, zinc-finger proteins could represent architectural factors, with implications for understanding genome organization.

Globally, CID structures exhibit low insulation scores in *Z. tritici*, consistent with previous observations in other fungi [32]. However, we identified a subset of highly insulated domains (HI-CIDs) that coincide with TE-rich, heterochromatic regions. Similar patterns have been described in *V. dahliae* and *Epichloë festucae*, where accessory genomic regions (AGRs) form structurally distinct and functionally specialized chromatin domains with stronger local interactions compared to surrounding regions [32,49]. In *Z. tritici*, these HI-CIDs also engage more frequently in strong interactions with one another compared to R-CIDs, which seems reminiscent of the spatial network formed by the “TAD cliques” observed in the human genome, where clusters of heterochromatic domains co-localize in 3D space [62]. Comparable chromatin bundling mechanisms have also been described in *C. elegans*, where H3K9me3 promotes chromosome compaction via HP1-mediated heterochromatin interactions [63]. In *N. crassa*, interspersed heterochromatin blocks are described to function as boundaries, structuring CIDs. Targeted removal of a heterochromatic block induces localized architectural changes with minimal effects on fitness [54]. The heterochromatin interaction network in *Z. tritici* shares conceptual similarities with the KNOT Engaged Elements (KNOTs) described in *Arabidopsis thaliana* [64]. KNOTs are heterochromatic islands within euchromatin that interact both within and between chromosomes. Notably, TEs preferentially insert into these regions, and under heat stress, KNOT-associated TEs are derepressed, demonstrating that these structures are dynamic and functional in TE regulation [65]. Previous studies in *Z. tritici* have shown that wheat infection is accompanied by a global activation of TEs and epigenetic changes [46]. We found that HI-CIDs are enriched for effector genes, and some of these CIDs are co-regulated *in planta,* suggesting potential de-repression of HI-CIDs during infection. While our study relies on *in vitro* Hi-C data, we recognize that host-induced chromatin remodeling likely plays a role in shaping 3D genome organization during plant infection. The low fungal biomass during early colonization stages currently limits the feasibility of generating *in planta* Hi-C maps in *Z. tritici*. Nonetheless, our finding that a subset of genes and TEs located within the same *in vitro*-defined CID tend to exhibit coordinated expression changes between axenic culture and infection suggests that CIDs may serve as functional domains of transcriptional co-regulation. Similar observations were drawn from studies in other fungal pathogens such as *V. dahliae* and *E. festucae*, where a subset of CIDs identified under axenic conditions showed regulatory relevance *in planta* [32,49]. These comparisons support the broader concept that core features of 3D genome architecture may be maintained across environmental conditions and can serve as predictive scaffolds for gene regulation. Future efforts to develop *in planta* approaches in fungal pathogens will be essential to directly validate this model and dissect the dynamics of chromatin folding during host colonization. Although the mechanisms and dynamics of chromatin folding during host colonization remain largely uncharacterized in fungi, the presence of specialized 3D genome organization represents an evolutionary strategy to compartmentalize potentially disruptive genomic elements while preserving their accessibility for regulated expression under specific developmental or stress-related conditions.

## CONCLUSION

The fundamental principles of genome folding, such as compartmentalization and domain organization, are broadly conserved across eukaryotes, even if the canonical definitions developed in metazoan systems do not always apply uniformly across the tree of life [55]. In the fungus *Z. tritici*, we uncovered a complex, multilayered chromatin architecture characterized by centromere clustering, spatial segregation between core and accessory chromosomes, a long-range heterochromatin network, and chromatin interacting domains. Together, these features likely contribute to this species’ remarkable genome plasticity, facilitating regulatory flexibility, adaptive potential, and pathogenicity. By generating a high-resolution map of *Z. tritici*’s 3D genome, we lay the groundwork for mechanistic investigations into how chromatin structure governs gene expression, transposable element dynamics, and genome evolution in filamentous fungi. More broadly, this work advances the emerging field of comparative 3D genomics in non-model species, opening new avenues for exploring genome architecture as a driver of fungal adaptation and plant-pathogen interactions.

## METHODS

### Sample preparation and Hi-C sequencing

The reference strain *Z. tritici* IPO323 [36] was preserved in 50% glycerol at −80°C. Pre-cultures were prepared with inoculations of ∼100uL of glycerol stock spore suspension onto solid YMS medium (4 g/L yeast extract, 4 g/L malt extract, 4 g/L sucrose, and 12 g/L of agar) and grown for 4 days at 18°C. Spores were scraped from plates and re-suspended into PBS buffer (8g/L NaCl, 1.44g/L Na_2_HPO_4,_ 0.2g/L KCl, 0.24g/L KH_2_PO_4_) for sample preparation for epigenomics sequencing as described in Möller et al. 2023 [41]. Spores were fixed with formaldehyde according to PhaseGenomics instructions (Seattle, Washington, USA). Hi-C experiments were performed with the Phase Genomics Proximo Hi-C Kit (Fungal), which combines four restriction enzymes to ensure homogeneous digestion at AT-rich genomic regions. The resulting libraries were sequenced at Biomarker Technologies GmbH (BMKgene, Münster, Germany). The sequencing was performed with Illumina NovaSeq X, paired-end 150bp reads. Sample preparation, Hi-C experiment, and sequencing were performed for two independent replicates.

### Hi-C data processing, binning, and visualization

Each Hi-C read-pairs was filtered and trimmed using Trimmomatic v0.39 [66] with the following settings ILLUMINACLIP:${adapters}:2:30:10 LEADING:15 TRAILING:15 SLIDINGWINDOW:5:15 MINLEN:50. We used HiCUP-0.9.2[67] to evaluate the quality of each Hi-C dataset. Reads were mapped against the reference genome assembly of *Z. tritici* IPO323 [36] using bwa-mem2 [68]. The construction and analysis of Hi-C interaction matrices were performed using the HiCExplorer suite of tools [60]. Each replicate interaction matrix was created using *hicBuildMatrix*, by aligning the paired reads to the reference genome digested *in silico* with the four restriction enzymes DpnII (GATC), DdeI (CTNAG), HinfI (GANTC), and MseI (TTAA). Contact matrices were individually corrected using the iterative correction and eigenvector decomposition (ICE) method via the *hicCorrectMatrix* function. The corrected contact matrices from the two biological replicates were combined to generate Hi-C matrices at 1 kb, 3 kb, 10 kb, and 50 kb. The Hi-C contact maps were generated with *hicPlotMatrix*. To identify the most significant chromatin interactions, we normalized the raw Hi-C contact matrices using the *hicTransform* from the HiCExplorer suite with the default --method obs_exp option. This method computes expected interaction frequencies as a function of genomic distance to account for the natural decay in contact frequency with increasing genomic separation. The most significant pairwise interactions were extracted by filtering the observed-over-expected values above 3.5 for inter- and 1.75 for intra-chromosomal interactions. The interaction link plots were generated using Circos-0.69 [69]. The HiCExplorer .h5 contact matrix files were converted into .cool format with *hicConvertFormat* for compatibility with the *cooltools* and *cooler* packages for visualization [70]. All scripts for the analyses and visualization are available at https://github.com/lorraincecile/Hi-C_Ztritici_IPO323 and <u>10.5281/zenodo.15395556</u>.

### Estimation of chromatin folding patterns with contact probability decay

To characterize chromatin folding behavior, we computed the decay of contact frequency as a function of genomic distance, a method grounded in polymer physics models of chromosome organization [4]. These models typically distinguish between different chromatin configurations based on the decay exponent (α) of the contact probability P(s), where *s* represents genomic separation. Contact decay slopes provide insights into the folding state of the chromatin (referred as “globule”): an exponent of ∼1.5 corresponds to an equilibrium globule, indicative of tightly knotted, inaccessible chromatin; values between 1 and 2 reflect a fractal globule, a compact yet unknotted state; and exponents < 1 are characteristic of a tension globule, normally associated with finer scale self-interacting domain units [71]. We used the *cooltools.expected_cis* [70] function to compute the average contact probability, P(s), separately for core and accessory chromosomes. Genomic distances and P(s) values were transformed to log10 scales, and a linear regression was fitted to the resulting log–log data to estimate the decay exponent (α), which serves as a proxy for the underlying chromatin folding state.

### Statistical Analysis of Chromosome Loss and Interaction Strength

To investigate the relationship between chromosome loss and interaction strength in *Z. tritici* IPO323, we extracted the average number of core-accessory chromosome strong interactions (observed vs. expected > 3.5), normalized by the size of the corresponding accessory chromosome. This normalization yielded the interaction strength per accessory chromosome. We compared this measure to the frequency of accessory chromosome loss following beta-tubulin inhibition, described in Habig et al. (2017) [38]. A log-transformed Poisson regression model was used to assess the impact of normalized interaction strength on accessory chromosome loss. The model was fitted using the glm function in R (version 4.4.1), with chromosome loss modeled as a Poisson-distributed response variable. The coefficients were tested for statistical significance using z-tests, with p-values calculated for both the intercept and the normalized interaction strength variable. Model diagnostics, including overdispersion testing using the *dispersiontest* function from the AER package [72], confirmed that the Poisson distribution was an appropriate fit. The fitted model was then used to predict chromosome loss, and the observed versus predicted values were visualized with ggplot2 [73].

### A/B compartmentalization analysis

Compartmentalization analysis used the matrix at 3 and 10 kb resolution with functions from the cooltools package [70]. We calculated PCA eigenvectors for intra- and inter-chromosomal contacts as *expected_cis* and *expected_trans,* respectively. Eigenvector values capture spatial segregation, with regions sharing similar values displaying more frequent contacts with one another than with regions showing dissimilar values. The intrachromosomal eigenvector profiles capture finer-scale chromatin features and reflects more local variability between contacts of the same chromosomes. GC content was used as a phasing track to orient the eigenvector values. The first three eigenvector values were manually checked, and the first eigenvector was determined to have the best compatibility with the interaction pattern of the contact maps. We used the 2.5% and 97.5% percentile eigenvector values as the minimum and maximum range to calculate the compartmentalization strength. The compartmentalization plot was generated using the *saddleplot* function [70].

### Chromatin interacting domains (CID) calling

We established 3 kb as the smallest bin size at which at least 1000 reads covered 80% of bins for each chromosome [9], and therefore the smallest scale at which we can reliably call CIDs (Supplementary Table S2). CID predictions were then generated using *hicFindTADs* for HiCExplorer [60] with default parameters using the contact matrix at 3kb bin size. This function calculates the insulation score by evaluating interaction frequencies between adjacent bins. CID boundaries were identified at significant local minima in physical interactions, where large differences between adjacent bins yield negative insulation scores, indicating “strong” insulation [60]. To assess the robustness of the predicted CIDs, we compared intra-domain interaction frequencies within annotated domains to those calculated in size-matched regions shifted upstream or downstream by half the length of each CID. All interaction values were ICE-normalized. We found that the original CIDs exhibited significantly higher intra-domain contact frequencies than both upstream- and downstream-shifted regions (Mann-Whitney U test; upstream: p-value < 0.05; downstream: p-value < 0.05), while the difference between upstream and downstream controls was not significant (p-value = 0.87) (Supplementary Fig. S12). CIDs were classified into “regular” (R-CIDs) and “high insulation” (HI-CIDs) CIDs by calculating the difference between each CID’s maximum insulation score and the minimum insulation scores of its adjacent CIDs. CIDs with a score difference within the 20th quantile, insulation scores above the median, and flanked by two neighboring CIDs with negative insulation scores were considered HI-CIDs.

### Analysis of genomic features

Gene annotations with associated GO terms, PFAM and effector predictions were sourced from Lapalu et al. (2023) [74] and accessible here (https://bioinfo.bioger.inrae.fr/portal/genome-portal/12/). Centromere detections and annotations were sourced from Schotanus et al. (2015), where centromere positions were experimentally determined using chromatin immunoprecipitation followed by sequencing (ChIP-seq) targeting the centromere-specific histone H3 variant CENH3 (CENP-A homolog) [39]. Core genes were identified as described in Minana-Posada et al. (2024)[75] as shared across 19 *Z. tritici* strains [47]. GO term enrichment analysis was performed using *topGO*’s *runGOEnrichment* function [76]. GO terms with no gene associations in the query lists or supported by only a single gene across the genome were excluded. False discovery rate (FDR) correction was applied to all p-values. Enrichment analyses of effectors, core genes, and PFAM domains within gene sets were performed using Fisher’s exact tests, with FDR correction applied to *p-*values. TE annotations were obtained from the manually curated library of Baril and Croll (2023) [77]. CID boundary motif identification and enrichment were performed with XSTREME with default parameters and compared to JASPAR database motifs [78] using Tomtom from the MEME suite tools [79]. We used the RIP composite index to identify RIP-like signatures in 1 kb sliding windows across the genome as described in Lorrain et al. (2021) [45]. A positive RIP composite index value reflects the presence of RIP-like mutations. The distribution of genes, TEs and RIP-like signatures across CIDs and their boundaries was analyzed using the EnrichedHeatmap package in R [80].

### Gene and TE expression analyses

RNA-seq data for *Z. tritici* IPO323 during *in planta* infection, comprising four distinct infection stages, was obtained from Haueisen et al. (2018) [81], while *in vitro* RNA-seq data was sourced from Möller et al. (2023) [41]. After trimming low-quality reads (ILLUMINACLIP:${adapters}:2:30:15 MINLEN:100), the resulting high-quality reads were aligned to the IPO323 reference genome using STAR [82], with parameters configured to allow the inclusion of multimapping reads (--outFilterMultimapNmax 200 and –winAnchorMultimapNmax 200). To capture the expression of genes and TEs, multi-mapper reads were reassigned using TEtranscript [83], which ensures precise allocation of reads to the transcript sequences [74] (https://bioinfo.bioger.inrae.fr/portal/genome-portal/12/) and TE library of *Z. tritici* IPO323 [77]. Differential expression analysis was performed using DESeq2 [84], comparing expression levels across the biotrophic phases (stages A and B), necrotrophic phases (stages C and D) [81], and *in vitro* growth conditions. To compare transcription between boundaries and CIDs, genes and TEs were analyzed separately, and statistical comparisons of expression levels and fold-change distributions were performed using two-sided Mann-Whitney U tests. The distribution of expression levels *in vitro* (log₂ RPKM) and differential expression comparing axenic and *in planta* conditions (log₂ fold-change), across CIDs and their boundaries, was analyzed using the EnrichedHeatmap package in R [80]. For each gene or TE, log₂-transformed values were calculated and centered on CID boundaries, extending 35 kb upstream and downstream. Heatmaps display normalized expression values per element, and the average signal profile across all regions is plotted above each heatmap. Absolute log₂ fold-change values were used to assess expression stability across conditions.

### Characterization of epigenetic profiles

Histone modifications, centromere detection, and cytosine DNA methylation (5mC) level enrichments across the *Z. tritici* IPO323 genome were assessed using publicly available chromatin immunoprecipitation followed by sequencing (ChIP-seq) and whole-genome bisulfite sequencing (WGBS) datasets. ChIP-seq datasets for CenH3 and H3K9me2 were obtained from Schotanus et al. (2015) [39], while datasets for H3K36me3, H4K20me3, H4K20me1, H3K27me3, H3K27me2, and H3K4me2 were sourced from Möller et al. (2023) [41]. Reads were preprocessed using Trimmomatic v0.39 [66], with parameters ILLUMINACLIP:${adapters}:2:30:10 LEADING:3 TRAILING:3 SLIDINGWINDOW:4:15 MINLEN:30 and ILLUMINACLIP:${adapters}:2:30:10:2:keepBothReads HEADCROP:5 LEADING:3 TRAILING:3 SLIDINGWINDOW:4:15 MINLEN:30 for the single and paired end samples respectively. The filtered reads were mapped to the IPO323 reference genome using Bowtie2 with default settings [85]. To account for the repetitive nature of TEs in the genome, multimapping reads were subsequently mapped using Allo [86], an approach tailored to improve mapping accuracy in repeat-rich regions. Reads with a mapping quality score below 30 and PCR duplicates identified by Picard v2.26.2 [87] were excluded from further analysis. To visualize enrichment levels across the genome, histone enrichment profiles were generated as bigWig and bedGraph files in bins of 10 bp, 3 kb, and 10 kb.

Bisulfite sequencing data was retrieved from Möller et al. (2021) [42] for cytosine DNA methylation analysis. Cultures were inoculated directly from glycerol stocks into liquid YMS medium and grown for 5 days. The Illumina HiSeq3000 paired-end raw reads were filtered using Trimmomatic [66] parameters HEADCROP:10 CROP:140 LEADING:3 TRAILING:3 SLIDINGWINDOW:4:15 MINLEN:50 and subsequently mapped and deduplicated using Bismark [88]. The methylated sites in all contexts (CG, CHG, CHH) were extracted using the function *bismark_methylation_extractor,* and filtered to remove sites not supported by at least four reads and more than 50% of the mapped reads. The weighted methylation average was calculated for 10 bp, 3 kb and 10 kb windows. The distribution of histone marks and 5mC levels across CIDs and their boundaries was analyzed using the EnrichedHeatmap package in R [80].

### Chromatin interacting domains co-expression analysis

To investigate the co-regulation of genes and TEs within CIDs, we constructed a linear model in Python as described in Winter et al. (2018) [49]. The model utilized the log2-fold change in gene expression between *in planta* and *in vitro* stages as the response variable, with CID membership as the main predictor. We developed nested models by incorporating additional genomic features known to affect gene expression, including GC content and distance to the nearest TE. The nested models’ performance was evaluated using the Akaike Information Criterion (AIC) and likelihood ratio tests. We defined CIDs as co-regulated CIDs with more than 10 co-expressed genes and TEs.

### Code availability

All custom scripts generated for data analyses and visualization are available at https://github.com/lorraincecile/Hi-C_Ztritici_IPO323. The Hi-C contact matrices and CID annotations as well as source data for figures and tables presented in this study are available at 10.5281/zenodo.15395556.

## Data availability

The Hi-C sequencing data is publicly available from the NCBI Gene Expression Omnibus (GEO) under the accession GSE296663. *In planta* [81] and *in vitro* [41] RNA-seq datasets are available at the GEO accession number GSE106136 and the accession number NCBI BioProjects PRJNA902413, respectively. The ChIP-seq datasets for CenH3 and H3K9me2 [39] are accessible through the SRA ID SRP059394. The ChIP-seq datasets for H3K36me3, H4K20me3, H4K20me1, H3K27me3, H3K27me2, and H3K4me2 [41] are available at the SRA ID SRP411885. Whole-genome bisulfite sequencing [42] data are available at SRA under BioProject ID PRJNA614493.

## Authors contributions

IG and CL designed the experiments. IG performed data acquisition. IG and CL performed the data analysis and interpretation. CL conceptualized the study. IG and CL wrote and revised the manuscript.

## Acknowledgments

We thank Prof. Bruce McDonald and ETH Zurich for funding the Hi-C sequencing. We thank Dr. Mareike Möller for her advice on performing Hi-C experiments in filamentous fungi.

## AI language model assistance

We used ChatGPT (developed by OpenAI) to refine and improve the grammar of the text. We reviewed and revised all outputs to ensure accuracy and alignment with the intended message.

## ETHICS DECLARATIONS

### Ethics approval and consent to participate

Not applicable.

### Consent for publication

Not applicable.

