## Supplementary Fig. for "Three-dimensional genome architecture connects chromatin structure and function in a major wheat pathogen"

Glavincheska & Lorrain

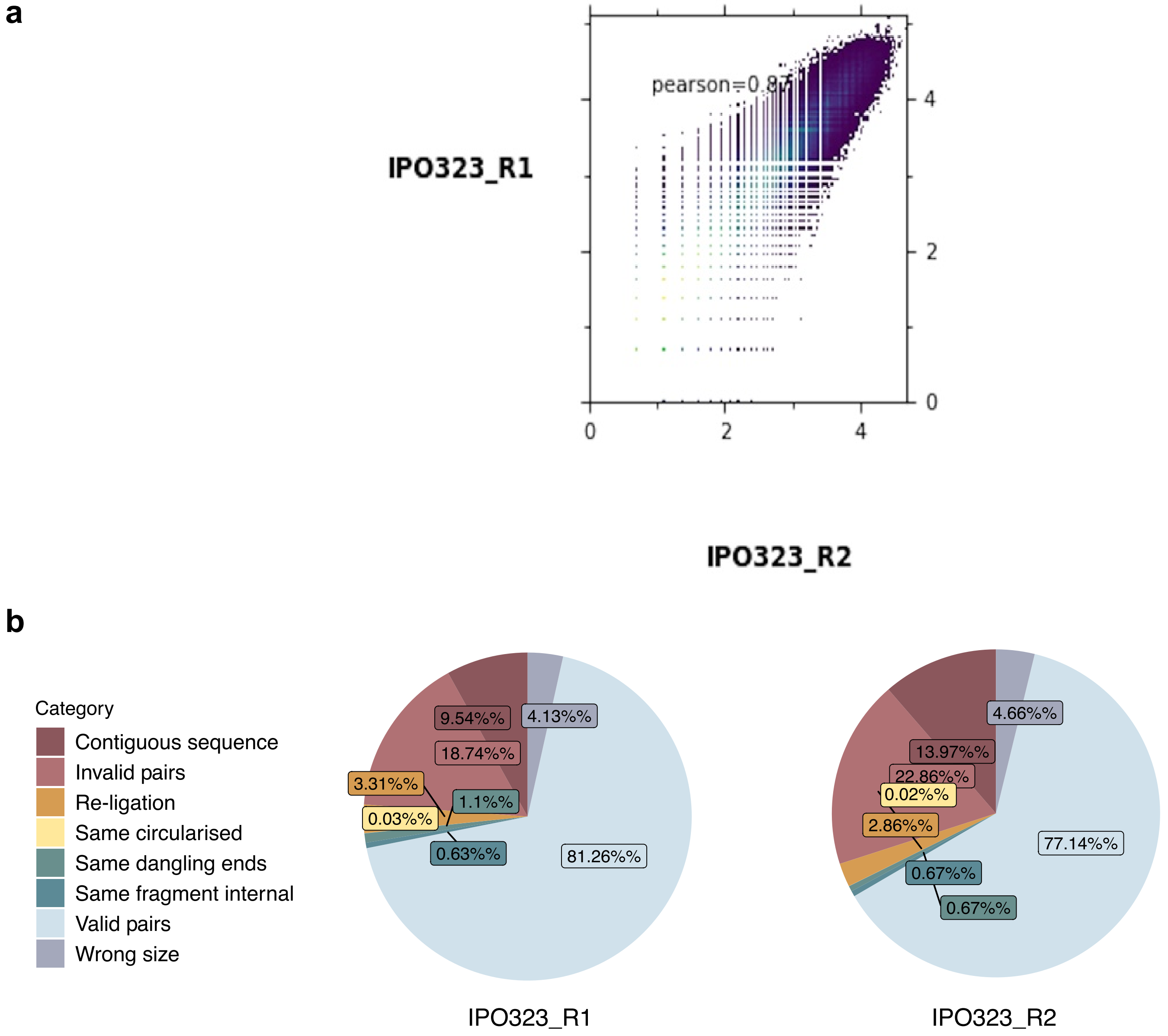


**Supplementary Fig. S1: Quality assessment of the Hi-C libraries replicates. a) Pearson correlation between Hi-C contact map replicates in *Zymoseptoria tritici* IPO323** (Pearson correlation = 0.87). b) Hi-C libraries quality filtering with the HiCUP-0.9.2 pipeline.

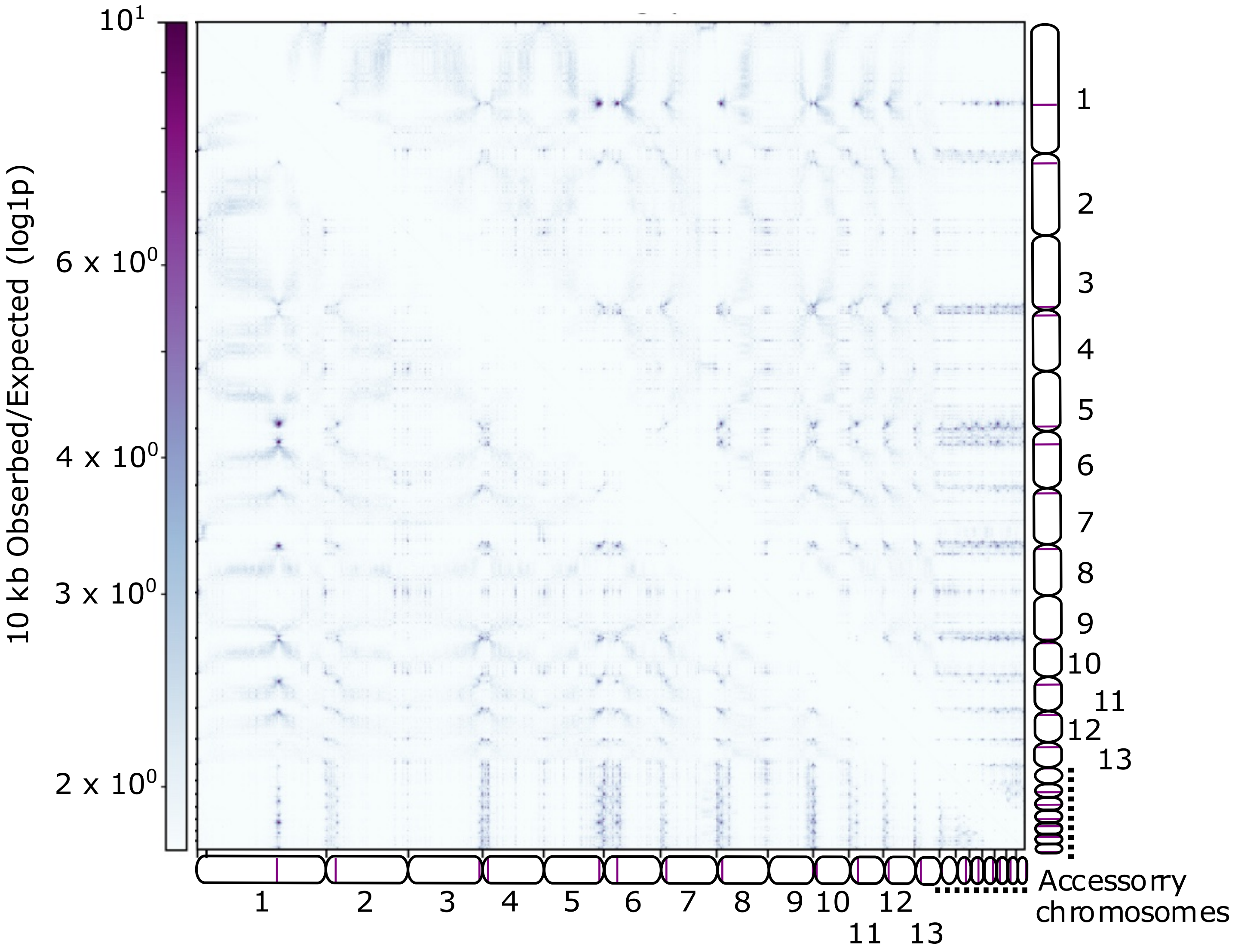


**Supplementary Fig. S2: *Zymoseptoria tritici* IPO323 Hi-C contact map 10kb bins normalized per genomic distance as observed/expected interactions.**

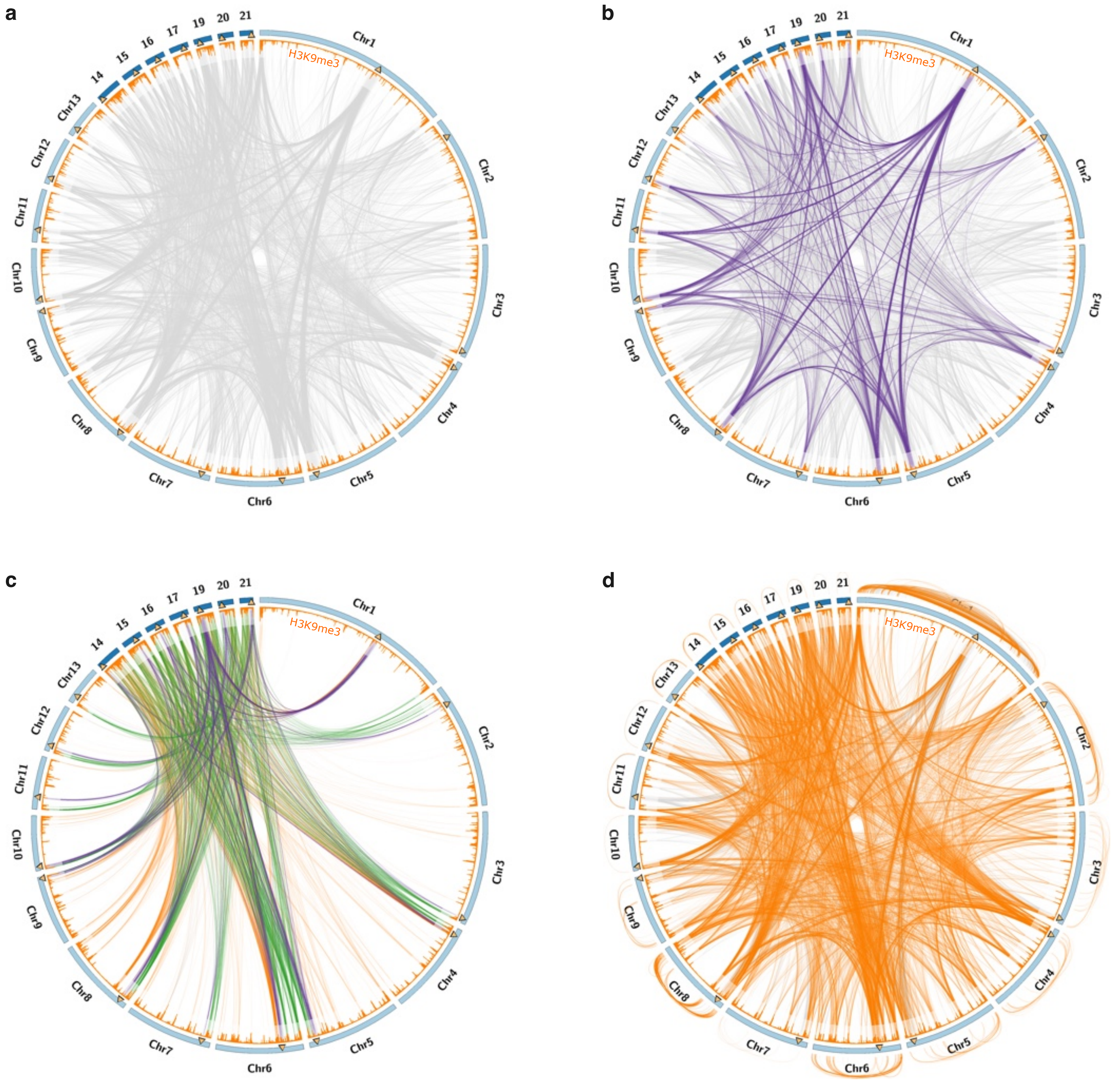


**Supplementary Fig. S3: Strongest interactions in *Zymoseptoria tritici* IPO323.** Links represent the strongest interactions with a log2 enrichment of observed-to-expected >3.5 at 10 kb bin resolution for a) all strong interactions, b) inter-chromosomal pericentromere-pericentromere interactions, c) accessory-core chromosome interactions with inter-chromosomal pericentromere-pericentromere interactions (purple), accessory-core chromosome (green), H3K9me3-H3K9me3 interactions (orange) or any other interactions (grey), d) H3K9me3-H3K9me3 interactions.

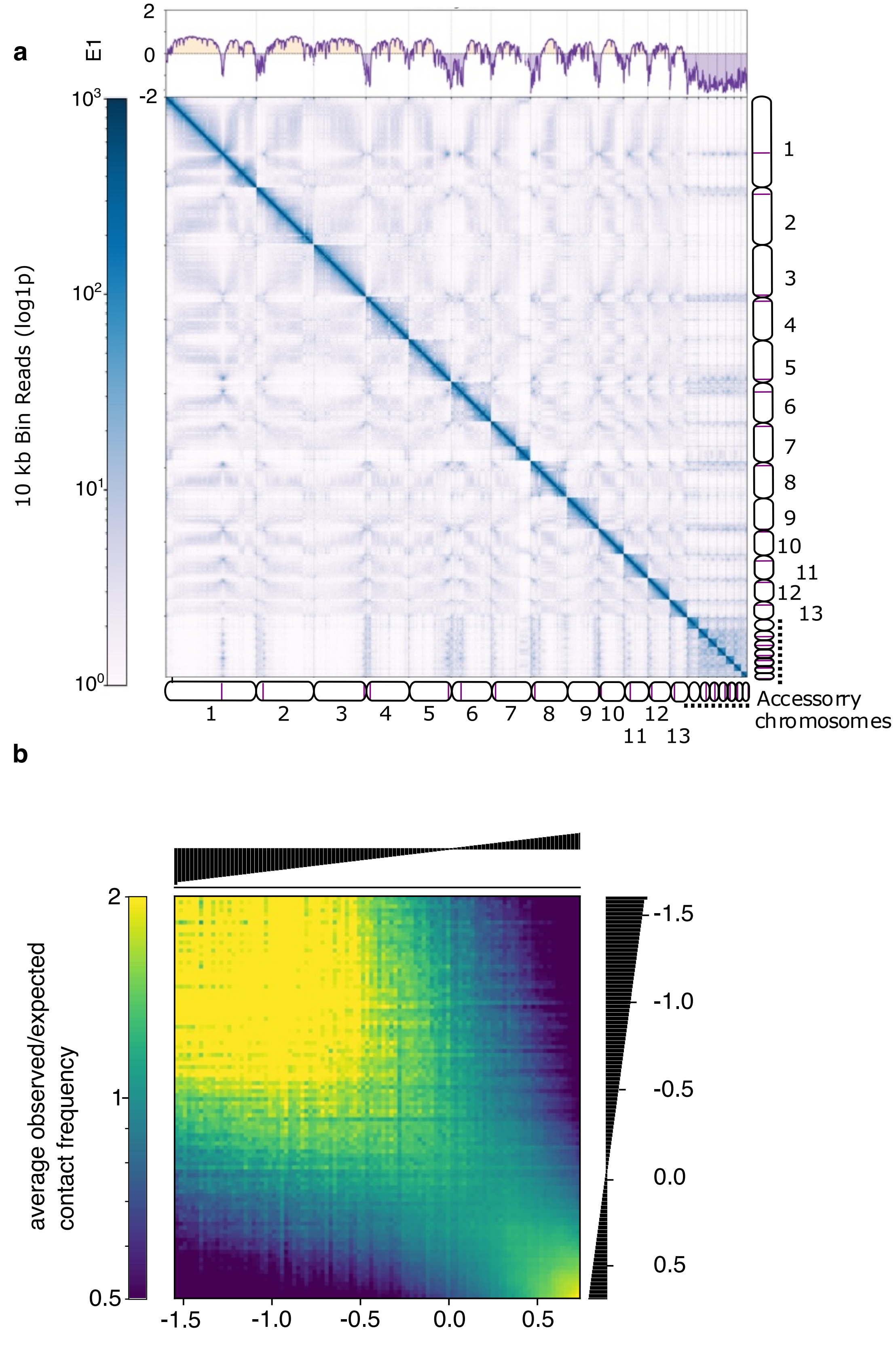


**Supplementary Fig. S4: Interchromosomal compartmentalization in *Zymoseptoria tritici* IPO323.** a) Self-interacting compartments were identified from Hi-C contact matrix at 10 kb resolution. Principal component analysis (PCA) of interchromosomal contact frequencies was used to extract the first eigenvector (E1), with positive and negative values annotated in yellow and purple, respectively. The eigenvector track is shown above each panel. The Hi-C contact map with 10 kb resolution is shown for all chromosomes, shown by ideograms on the side, with centromeres represented in purple. b) Saddle plot representing genome-wide Hi-C interactions at 10 kb resolution. Homotypic interactions between regions associated with a negative E1 are represented in the top-left, while positive E1 region interactions are observed in the bottom-right.

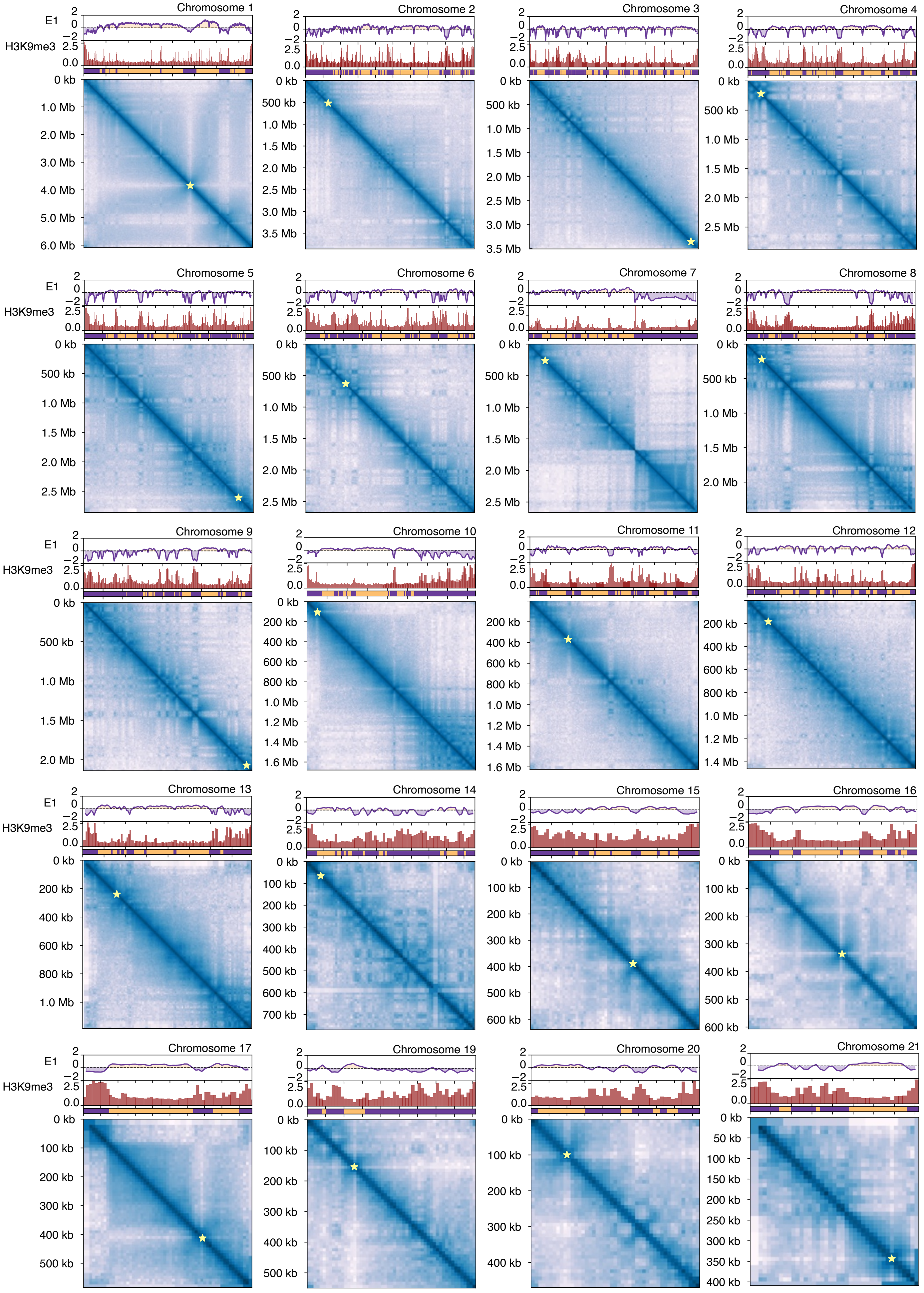


**Supplementary Fig. S5: Intrachromosomal A/B compartments in *Zymoseptoria tritici* IPO323.** For each chromosome, A/B compartments were identified from Hi-C contact matrices at 10 kb resolution. Principal component analysis (PCA) of intrachromosomal contact frequencies was used to extract the first eigenvector (E1), with positive and negative values corresponding to A (yellow) and B (purple) compartments, respectively. The eigenvector track is shown above each panel. Enrichment of H3K9me3 per 10 kb window is displayed as a histogram below the eigenvector track. Hi-C contact maps are shown for individual chromosomes. Centromeres are represented by yellow stars.

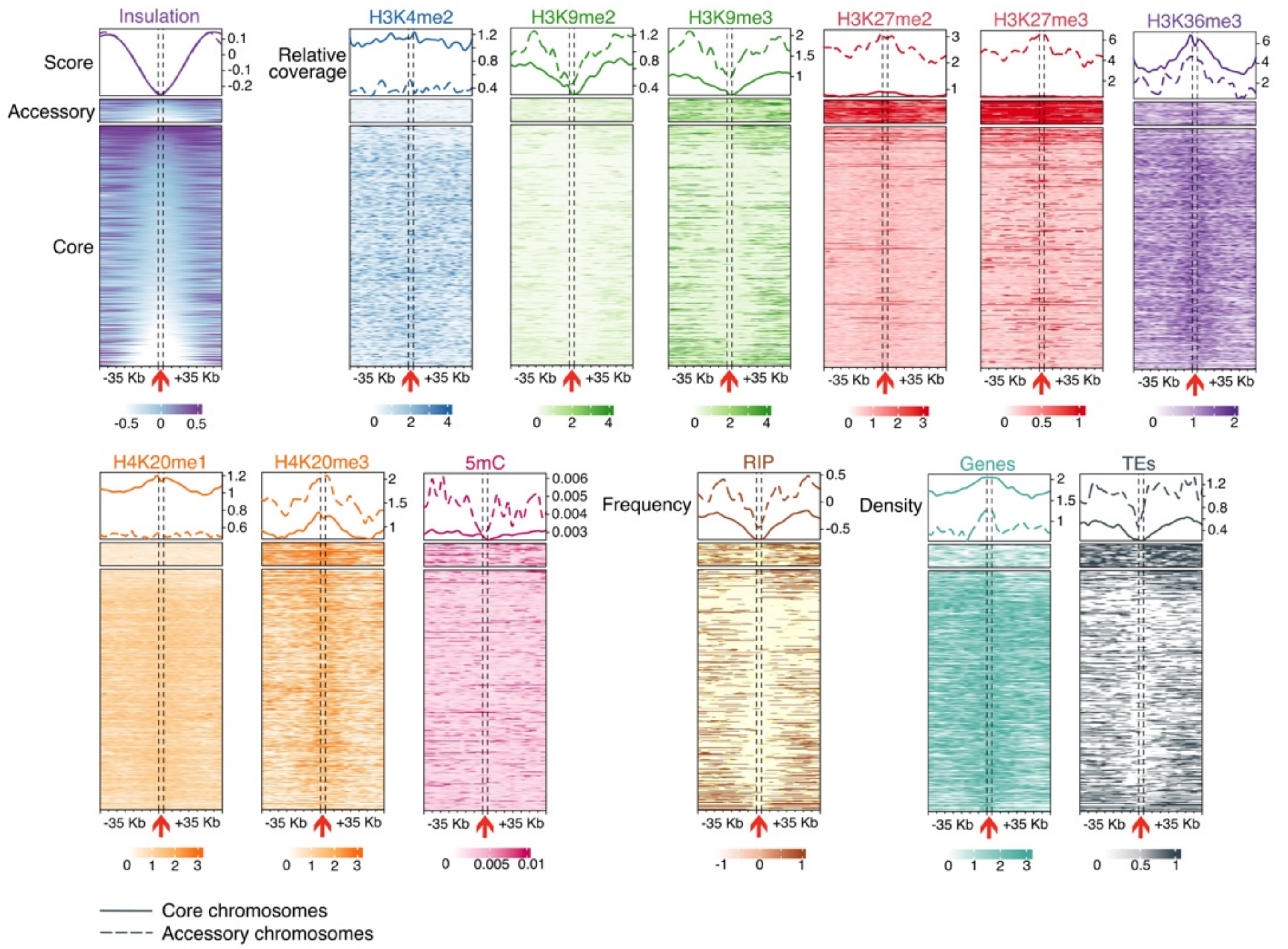


**Supplementary Fig. S6: Chromatin interacting domain (CID) boundaries characteristics in core and accessory chromosomes of *Zymoseptoria tritici* IPO323.** Heatmaps of chromatin insulation scores, enrichment in histone modifications (H3K4me2, H3K9me2, H3K9me3, H3K27me2, H3K27me3, H3K36me3, H4K20me1, H4K20me3), 5-methylcytosine DNA methylation (5mC), repeat-induced point mutation (RIP) composite index frequency, genes, and TEs density centered on domain boundaries, with 35 kb upstream and downstream regions shown per row. The top panel shows the average profile across all core (solid lines) and accessory (dash lines) chromosomes boundaries depicted in the heatmap.

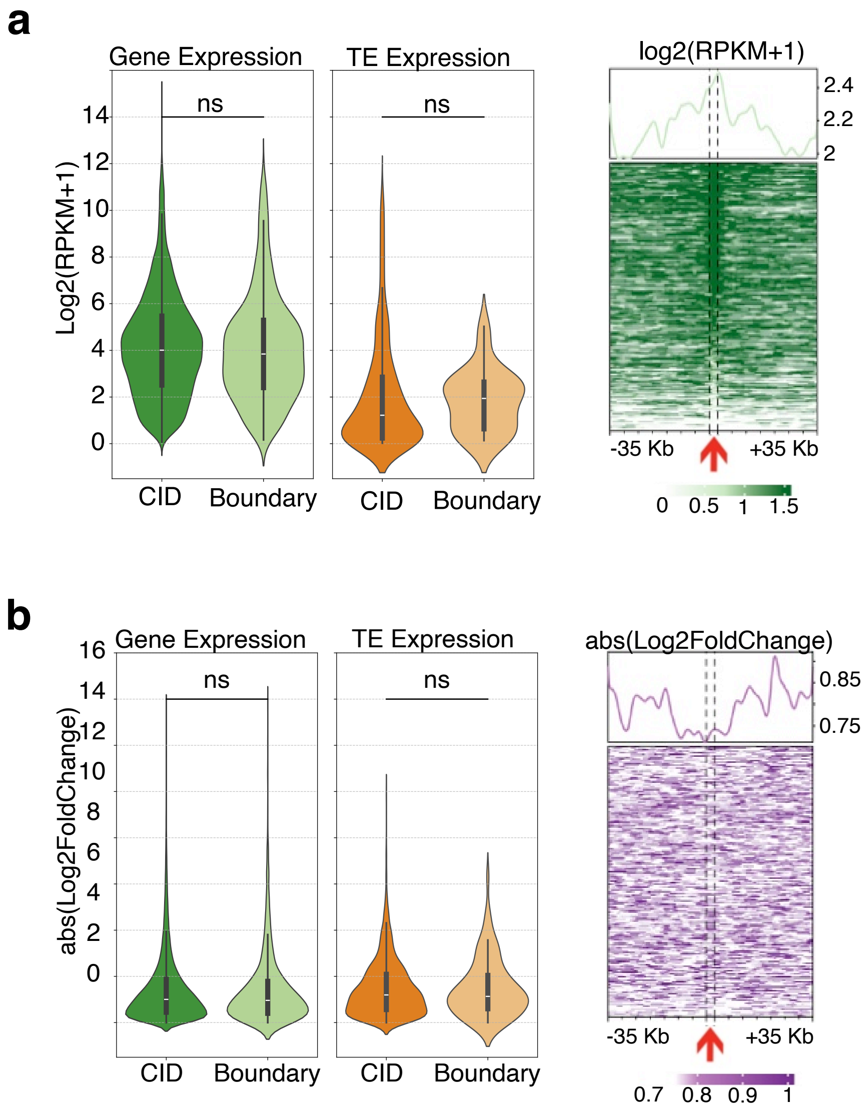


**Supplementary Fig. S7: In vitro expression levels of genes and transposable elements in chromatin interacting domains (CIDs) and boundaries.** a) Violin plots (left) show log$_2$-transformed expression levels (log2 RPKM+1) for genes (green) and TEs (orange) located within CIDs or at CID boundaries under axenic culture conditions. The heatmap (right) shows normalized gene expression centered on CID boundaries, with 35 kb upstream and downstream per row. The top panel displays the average log$_2$(RPKM+1) profile across all boundaries. b) Violin plots (left) show the absolute log$_2$ fold-change in gene and TE expression (infection vs. axenic) for elements within CIDs or at boundaries. The heatmap (right) shows absolute expression changes centered on CID boundaries (±35 kb), with the top panel showing the average profile. Statistical significance of differences between CIDs and boundaries was assessed using a two-sided Mann-Whitney U test with asterisks indicating significance levels (ns: non-significant).

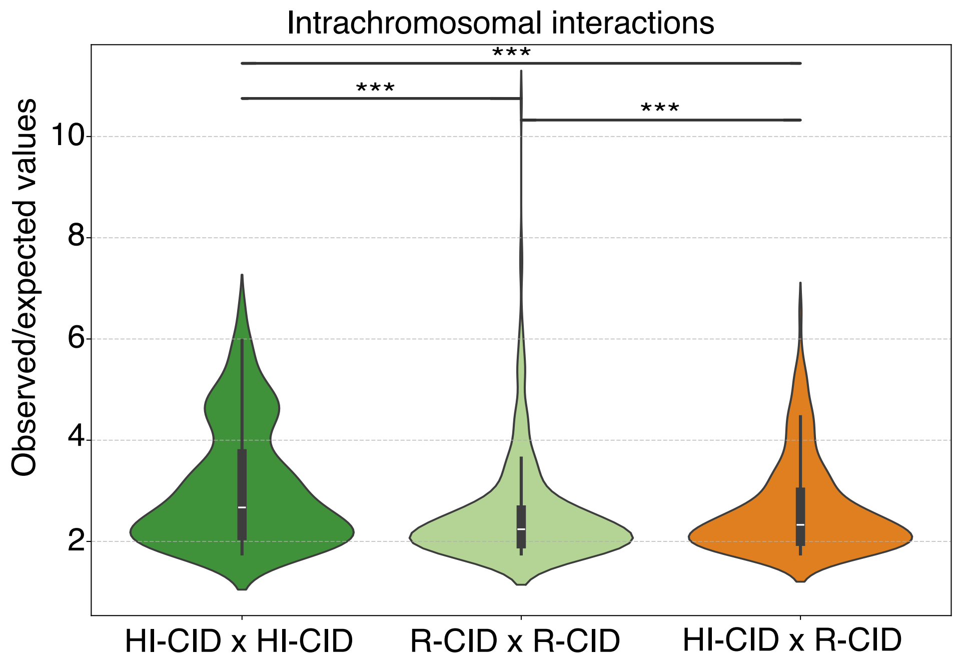


**Supplementary Fig. S8: Strongest interactions between chromatin interacting domain (CIDs).** Distribution of observed-to-expected contact ratios > 1.75 from the 10 kb resolution interaction matrix, highlighting the strongest intrachromosomal interactions classified as either highly insulated CIDs (HI-CIDs) or regular CIDs (R-CIDs). Statistical significance of differences between groups was assessed using a two-sided Mann-Whitney U test with asterisks indicating significance levels (ns: non-significant; *: p-value < 0.05; **: p-value < 0.01; ***: p-value < 0.001).

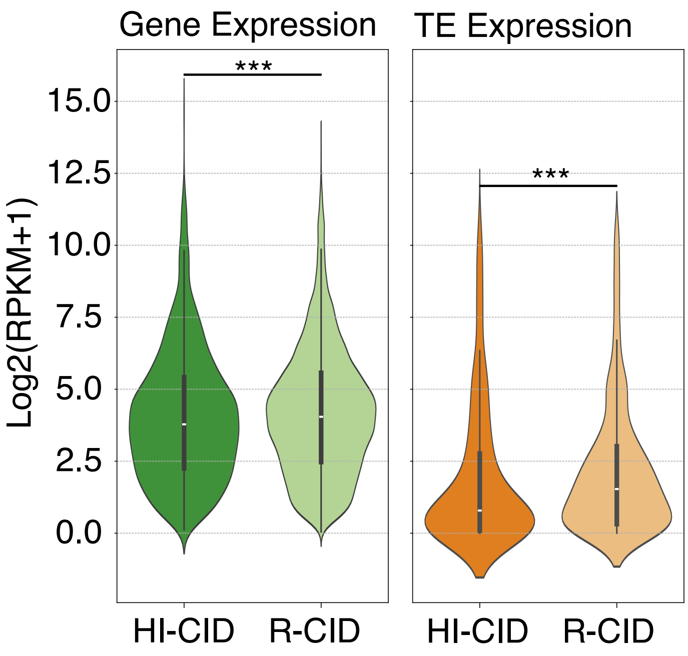


**Supplementary Fig. S9: In vitro Expression levels of genes in regular chromatin interacting domains (R-CIDs) and highly insulated chromatin interacting domains (HI-CIDs).** Genes are represented in green and transposable elements in orange. Statistical significance of differences between HI-CIDs and R-CIDs was assessed using a two-sided Mann-Whitney U test with asterisks indicating significance levels (ns: non-significant; *: p-value < 0.05; **: p-value < 0.01; ***: p-value < 0.001).

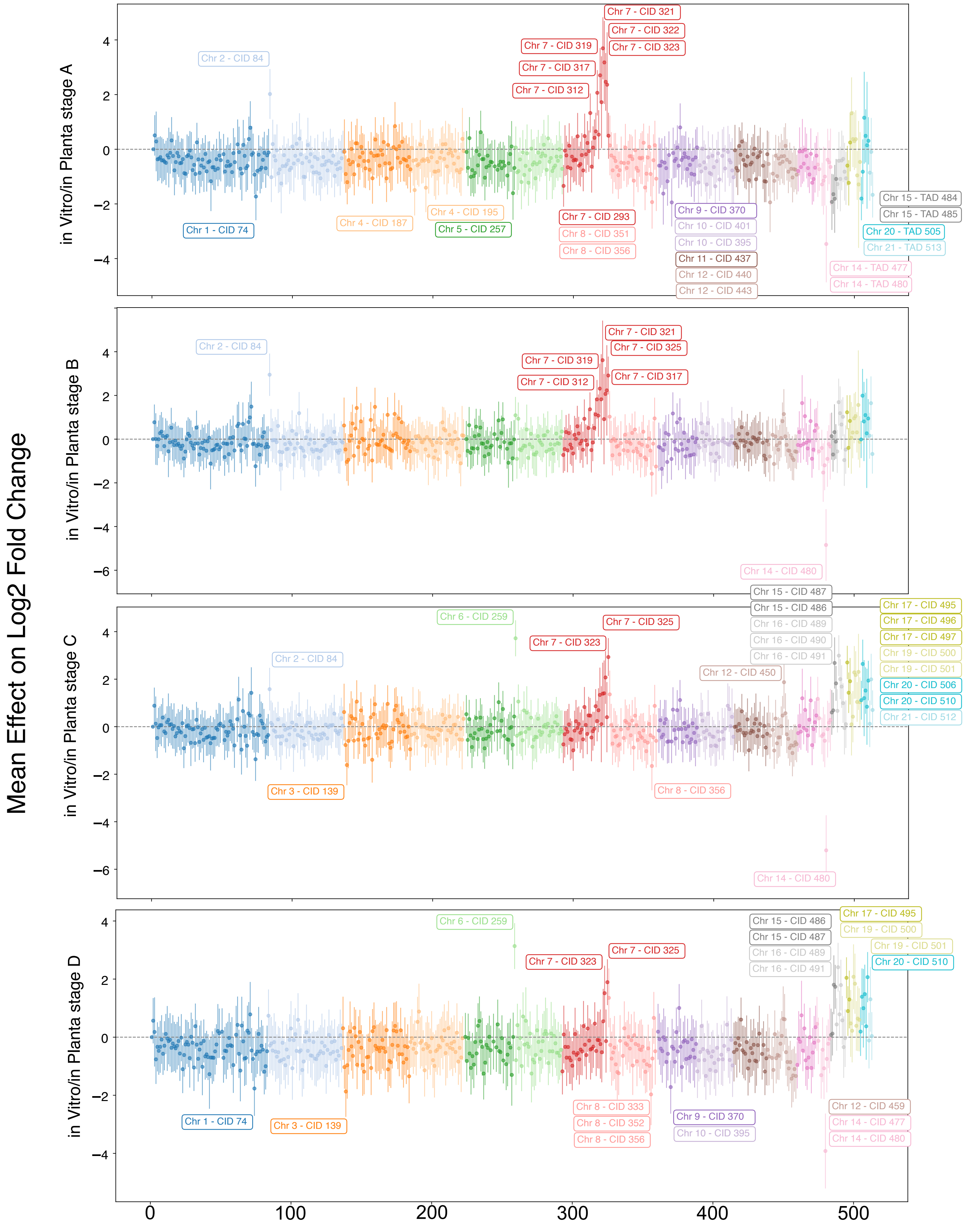


**Supplementary Fig. S10: Co-regulated chromatin interacting domains (CIDs) during wheat infection stages.** Effect sizes from linear regression models quantified the contribution of each topologically associating domain (CID) to differential gene expression between in vitro growth on YMA medium and various in planta infection stages (stage A: early biotrophic, stage B: late biotrophic, stage C: early necrotrophic, and stage D: late necrotrophic). Each point represents the mean effect size per CID, accompanied by a 95% confidence interval. CIDs with statistically significant effects (i.e., confidence intervals not overlapping zero) are highlighted in color and annotated with their chromosome and CID identifiers. Values reflect normalized means derived from two independent biological replicates [81].

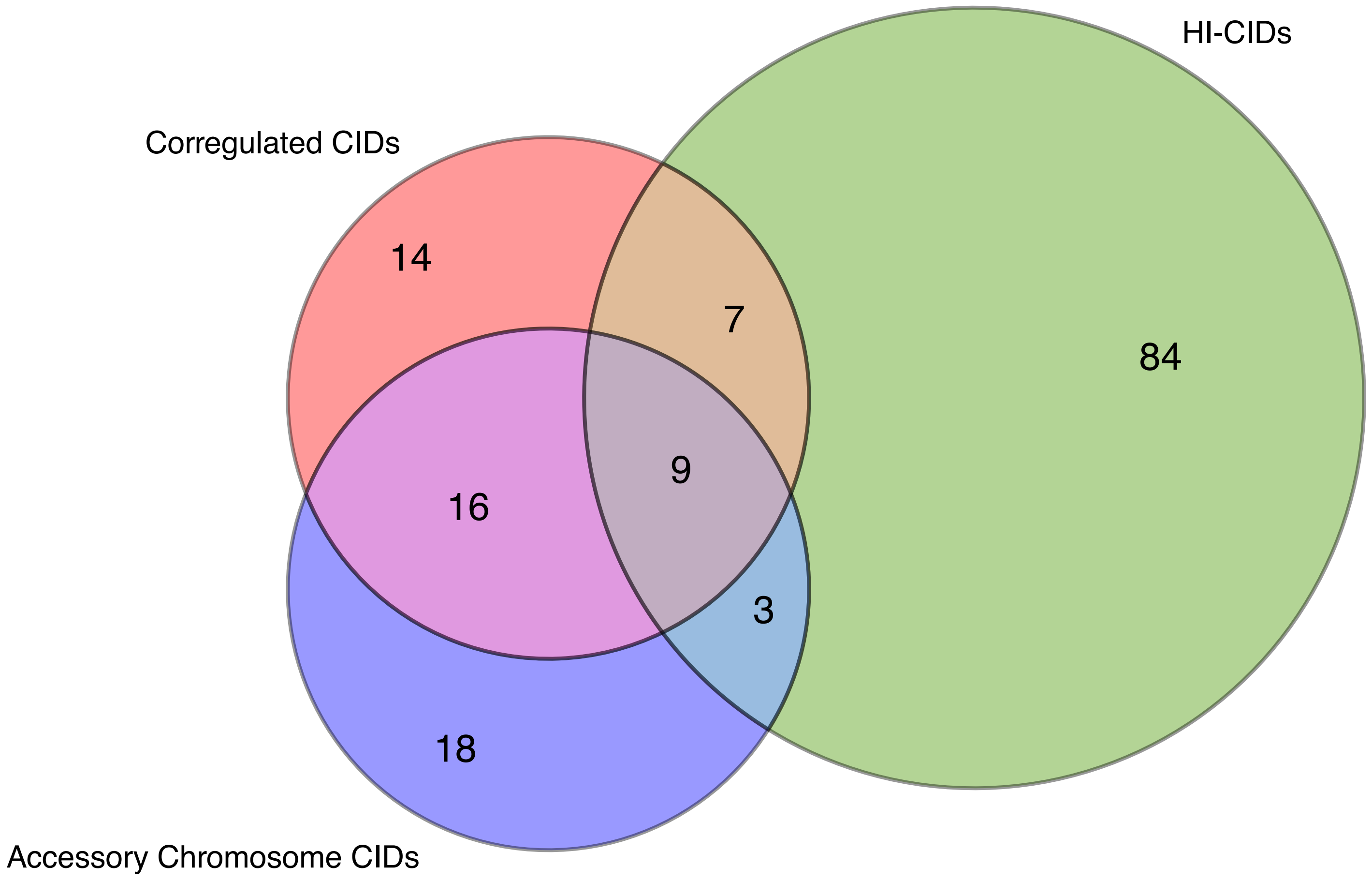


**Supplementary Fig. S11: Overlap between chromatin interacting domain (CID) categories**. Venn diagram illustrating the overlap among three categories of topology-associated domains (CIDs): co-regulated CIDs (defined as containing  10 genes or transposable elements with coordinated expression), highly insulated CIDs (HI-CIDs; identified by elevated insulation scores), and accessory chromosome CIDs (located on accessory chromosomes).

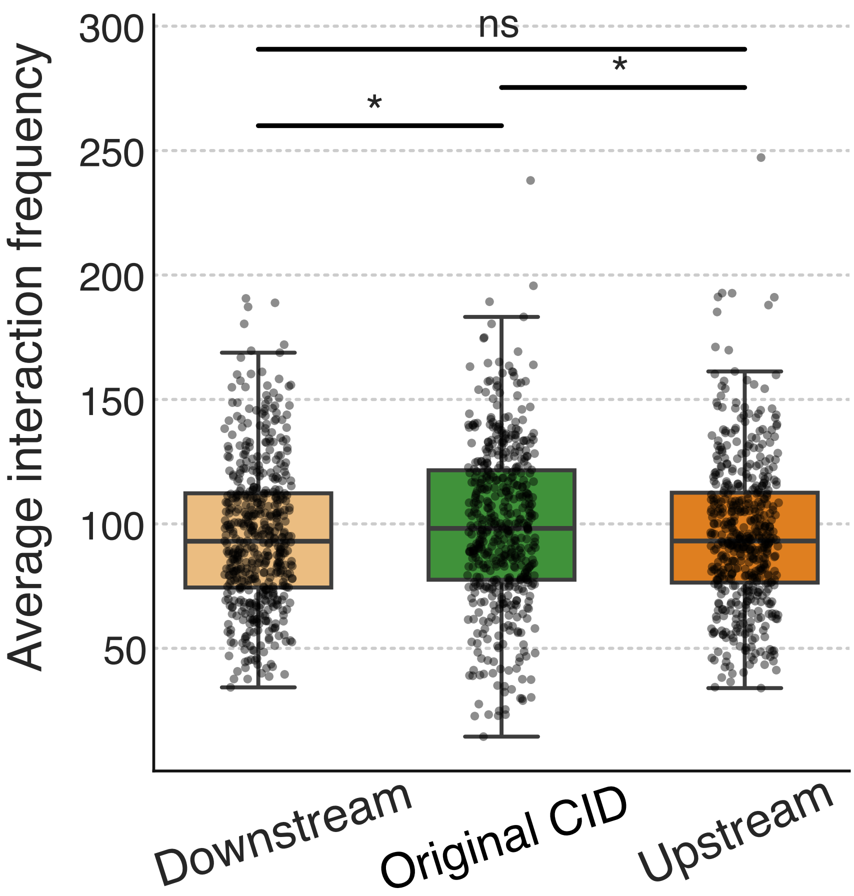


**Supplementary Fig. S12: Validation of CID boundary predictions based on intra-domain interaction frequencies.** Boxplots show the average intra-domain interaction frequency for annotated CIDs ("Original CID") compared to control regions of identical size shifted upstream or downstream by half the domain length (two-sided Mann-Whitney U test with asterisks indicating significance levels (ns: non-significant; *: p-value < 0.05).
